# Mitotically heritable, RNA polymerase II-independent H3K4 dimethylation stimulates *INO1* transcriptional memory

**DOI:** 10.1101/2022.02.11.480054

**Authors:** Bethany Sump, Donna Garvey Brickner, Agustina D’Urso, Seo Hyun Kim, Jason H. Brickner

**Affiliations:** Department of Molecular Biosciences, Northwestern University, Evanston, Illinois, USA

## Abstract

For some inducible genes, the rate and molecular mechanism of transcriptional activation depends on the prior experiences of the cell. This phenomenon, called *epigenetic transcriptional memory*, accelerates reactivation and requires both changes in chromatin structure and recruitment of poised RNA Polymerase II (RNAPII) to the promoter. Memory of inositol starvation in budding yeast involves a positive feedback loop between transcription factor-dependent interaction with the nuclear pore complex and histone H3 lysine 4 dimethylation (H3K4me2). While H3K4me2 is essential for recruitment of RNAPII and faster reactivation, RNAPII is not required for H3K4me2. Unlike RNAPII-dependent H3K4me2 associated with transcription, RNAPII-independent H3K4me2 requires Nup100, SET3C, the Leo1 subunit of the Paf1 complex and, upon degradation of an essential transcription factor, is inherited through multiple cell cycles. The writer of this mark (COMPASS) physically interacts with the potential reader (SET3C), suggesting a molecular mechanism for the spreading and re-incorporation of H3K4me2 following DNA replication.

## INTRODUCTION

Cells react to changes in their environment by altering gene expression, primarily by regulating transcription of inducible genes. The rate of induction of such genes is a product of enhancer and promoter activity (ngoc et al., 2017), but can also be influenced by the previous experiences of the cells. A number of genes from yeast, flies, worms, mammals, and plants are more strongly induced following a previous exposure to a stimulus and this primed state can persist for 4-14 mitotic cell divisions (Brickner et al., 2007; D’Urso et al., 2016; Gialitakis et al., 2010; Lämke and Bäurle, 2017; Light et al., 2013, 2010; Maxwell et al., 2014; Pascual-Garcia et al., 2017; Siwek et al., 2020; Sood and Brickner, 2017). These phenomena, collectively called epigenetic transcriptional memory, quantitatively alter gene expression in response to recent experiences (D’Urso and Brickner, 2017, 2014). Because only a subset of genes induced under a particular condition exhibits memory (Light et al., 2013), this can also result in a qualitative change in the global expression pattern.

Although transcriptional memory in different organisms or genes has unique features, several common, conserved mechanisms have been identified. For example, in budding yeast, flies and mammals, transcriptional memory requires the nuclear pore protein Nup98 (homologous to Nup100 in yeast). This protein physically interacts with the promoters of genes that exhibit memory and loss of Nup98/Nup100 disrupts memory (Light et al., 2013, 2010; Pascual-Garcia et al., 2017). The interaction with Nup98 with chromatin in flies and mammals can occur away from the nuclear pore complex (NPC; Capelson et al., 2010; Kalverda et al., 2010). However, during memory, the interaction of promoters with Nup98/Nup100 in both yeast and flies occurs at the pore.

Transcriptional memory is associated with local changes in chromatin modifications and chromosome folding. In *Drosophila*, ecdysone memory involves a long-distance promoter-enhancer interaction that is induced by ecdysone and is strengthened/stabilized by Nup98 (Pascual-Garcia et al., 2017). In yeast, plants and mammals, histone modifications of the promoter are required for memory: dimethylation of histone H3 lysine 4 (H3K4me2) is generally associated with memory in yeast and mammals (D’Urso et al., 2016; Gialitakis et al., 2010; Light et al., 2013, 2010), while in plants, several forms of epigenetic memory are associated with either H3K4me2 or H3K4me3 (Lämke and Bäurle, 2017). In flies and mammals, Nup98 physically interacts with the H3K4 methyltransferases Trx and Set1A/COMPASS, respectively (Franks et al., 2017; Pascual-Garcia et al., 2014). Nup98/Nup100 is required for memory-associated H3K4me2 in yeast and mammals and conditional inactivation of either COMPASS (the methyltransferase) or SET3C (a histone deacetylase complex that binds H3K4me2; Kim and Buratowski, 2009) leads to rapid loss of memory in yeast (D’Urso et al., 2016; Light et al., 2013, 2010). Importantly, the dimethylation of histone H3 lysine 4 during memory in yeast is carried out by a form of COMPASS that lacks the Spp1 subunit (Spp1^-^ COMPASS), preventing trimethylation (D’Urso et al., 2016).

Finally, transcriptional memory in yeast and mammals leads to binding of a poised RNA polymerase II pre-initiation complex (RNAPII PIC; D’Urso et al., 2016; Light et al., 2013, 2010). A similar phenomenon, called RNA polymerase II “docking” has been reported in *C. elegans* (Maxwell et al., 2014). Work from yeast suggests that this poised RNAPII PIC is distinct from active RNAPII PIC in two ways. First, it fails to recruit Cdk7 (Kin28 in budding yeast), the kinase that phosphorylates serine 5 on the RNAPII carboxy terminal domain upon initiation (D’Urso et al., 2016). Second, it remains associated with Mediator kinase Cdk8 (Ssn3 in budding yeast; D’Urso et al., 2016). Cdk8 is also associated with poised promoters in HeLa cells (D’Urso et al., 2016), and has been found to regulate initiation (Akoulitchev et al., 2000; Pavri et al., 2005) suggesting that it plays a conserved role in transcriptional poising. Conditional depletion of yeast Ssn3 from the nucleus leads to loss of memory (D’Urso et al., 2016). Therefore, transcriptional memory is a conserved phenomenon that may involve a conserved core mechanism.

The highly inducible budding yeast gene *INO1* has served as a model for epigenetic transcriptional memory. *INO1* is an essential enzyme that catalyzes the conversion of glucose to *myo*-inositol for the biosynthesis of phosphatidylinositol. Our current understanding of *INO1* memory is summarized in Figure 1A. Repressed *INO1* localizes in the nucleoplasm. Upon activation, the gene repositions to the nuclear periphery through interaction with the NPC (Ahmed et al., 2010; Brickner and Walter, 2004; Sumner et al., 2021). Interaction of active *INO1* with the NPC requires two transcription factors, Put3 and Cbf1, that bind to the upstream DNA zip codes GRSI and GRSII (Figure 1A; Ahmed et al., 2010; Brickner and Walter, 2004; Randise-Hinchliff et al., 2016). Upon repression, recently-repressed *INO1* acquires memory-specific chromatin marks, poised RNAPII PIC and associates with the NPC by another mechanism (Brickner et al., 2007; Light et al., 2013, 2010). Whereas nucleosomes in the promoter and 5’ end of active *INO1* are hyper-acetylated and show both H3K4me2 and H3K4me3, the nucleosomes at the 5’ end of recently-repressed *INO1* are hypo-acetylated and show only H3K4me2 (D’Urso et al., 2016; Light et al., 2013). Also, the histone variant H2A.Z is incorporated into an upstream nucleosome during *INO1* memory (Brickner et al., 2007; Light et al., 2010). The interaction with the NPC requires the Sfl1 transcription factor, the Memory Recruitment Sequence (MRS) DNA zip code, to which Sfl1 binds, and the nuclear pore protein Nup100 (D’Urso et al., 2016; Light et al., 2013, 2010). Inactivation of any of these factors disrupts memory without affecting repression or activation. *INO1* memory, as monitored by gene positioning, chromatin changes or histone modification, is maintained in mother cells and inherited to daughter cells through up to four mitotic divisions before being lost (Brickner et al., 2007; D’Urso et al., 2016).

**Figure 1.**
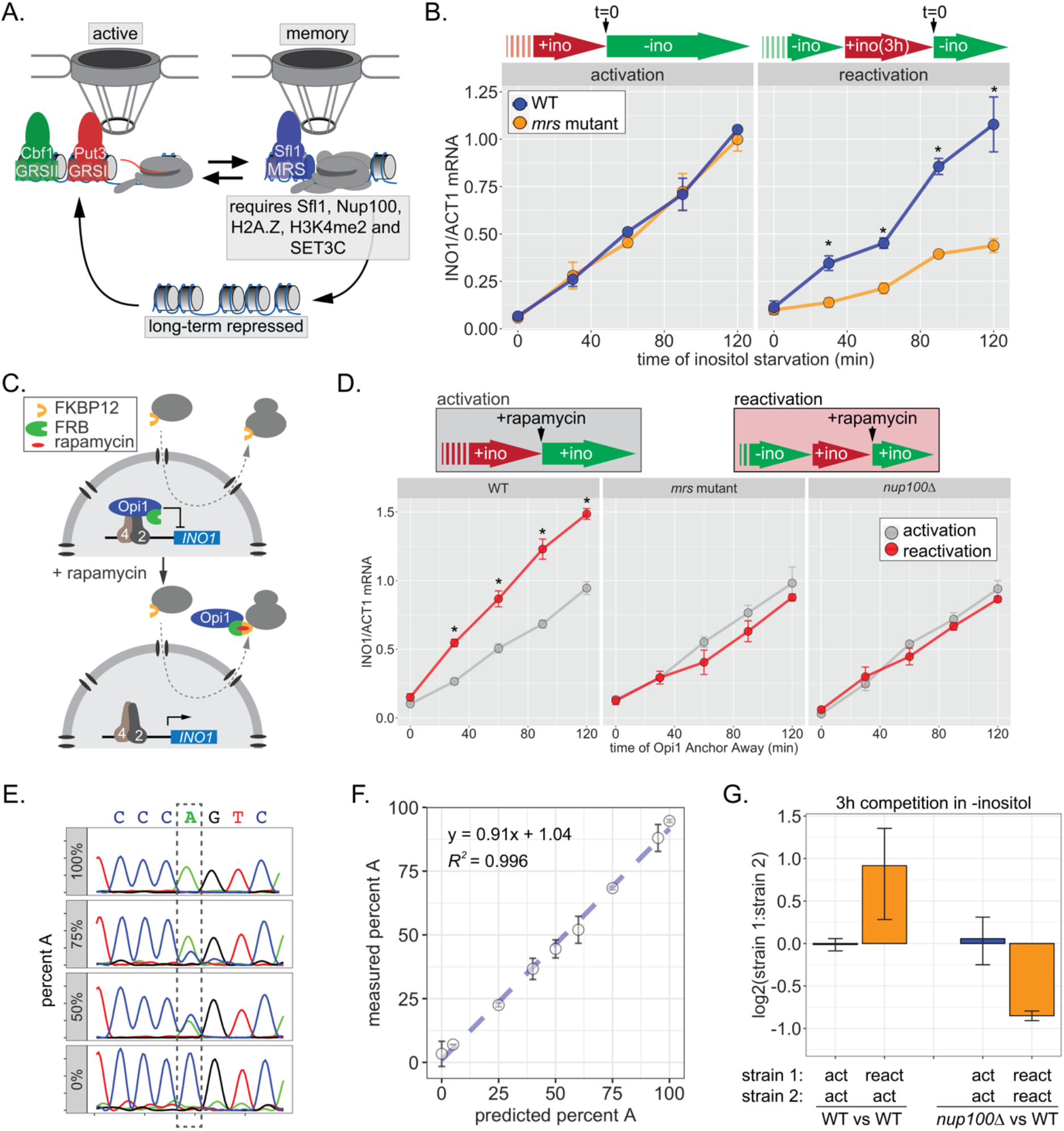
*INO1* transcriptional memory stimulates faster transcription and provides a competitive fitness advantage. **(A)** Model of *INO1* in the active, memory, and long-term repressed states, highlighting factors that are specifically required for memory. **(B)** Activation (left) and reactivation (right) of *INO1* in wild type and *mrs* mutant strains upon starvation of inositol. Cells were harvested at indicated time points and the *INO1* mRNA was quantified relative to *ACT1* mRNA by RT-qPCR (*p-value < 0.05 from one-tailed t-test comparing WT and *mrs* mutant, alternative = greater). **(C)** Schematic of Anchor Away of Opi1 to induce *INO1*. **(D)** Top: experimental scheme for synthetic activation and reactivation of *INO1*. Activation and reactivation of *INO1* in wild type (left), *mrs* mutant (middle) and *nup100Δ* (right) strains upon removal of Opi1 by Anchor Away. Bottom: Cells were harvested at indicated time points and *INO1* mRNA was quantified relative to *ACT1* mRNA by RT-qPCR (*p-value < 0.05 from one-tailed t-test comparing reactivation and activation, alternative = greater). **(E)** Chromatograms resulting from sequencing mixtures of strains having either “A” or “C” SNP within an integrated plasmid, as indicated (dashed box). **(F)** Standard curve comparing the predicted percentage of strain A (as estimated by O.D._600_) with the measured percentage of A (as quantified by the relative area under the peaks, as shown in **E**). **(G)** Relative fitness of competing strains, as indicated. The log_2_ ratio of the abundance of the two strains after 3 hours of competition in media lacking inositol is shown. For panels B, D, F and G, data are averages of 3 biological replicates ± standard error of the mean (SEM).

Here we exploited our knowledge of *INO1* transcriptional memory to address four critical questions. First, although transcription rates are impacted by memory, how it impacts fitness has not been generally assessed. We find that *INO1* memory provides a competitive fitness advantage during inositol starvation that is Nup100-dependent. Second, we assessed the function of H3K4me2 during memory. Sfl1 is required for H3K4 dimethylation and H3K4me2 is essential for both Sfl1 binding and for RNAPII recruitment during *INO1* memory, suggesting that memory involves a chromatin-dependent positive feedback loop. Third, we define ways in which the dimethylation of H3K4 during transcription is different from dimethylation of H3K4 during memory. While transcription-associated H3K4 methylation is RNAPII-dependent, memory-associated H3K4me2 is RNAPII-independent and requires both overlapping and distinct factors. Finally, we explore the molecular mechanism of epigenetic inheritance of *INO1* memory. We rule out that protein production during inositol starvation promotes memory and instead, highlight the critical role of heritable H3K4me2. Although establishing H3K4me2 during memory requires Sfl1, once established, H3K4me2 is maintained and inherited ∼4 cell divisions in the absence of Sfl1. A putative reader of H3K4me2 (SET3C) physically interacts with the writer of H3K4me2 (COMPASS), suggesting a molecular mechanism by which H3K4me2 is inherited during DNA replication. This work provides a compelling example of a heritable histone modification that stimulates future transcription and is inherited over a shorter timescale than other heritable histone modifications like H3K9 methylation or H3K27 methylation.

## RESULTS

### *INO1* epigenetic transcriptional memory stimulates faster transcription and promotes competitive fitness

Factors that are specifically required for *INO1* memory stimulate faster reactivation. For example, while the rate of *INO1* activation (+inositol ⟶ -inositol) is unaffected by mutations in the MRS, the rate of *INO1* reactivation (-inositol ⟶ +inositol, 3h ⟶ -inositol) is clearly decreased by such mutations (Figure 1B). Thus, Sfl1 binding to the MRS is important for enhancing the rate of *INO1* reactivation and has no role in *INO1* activation. However, this experiment highlights a paradox: although poised RNAPII PIC is associated with the recently-repressed *INO1* promoter prior to reactivation (D’Urso et al., 2016; Light et al., 2010), the rate of reactivation is not obviously faster than the rate of activation (Figure 1B). This has led to the suggestion that *INO1* does not exhibit transcriptional memory (Halley et al., 2010). However, it is possible that the rate at which inositol starvation is *perceived* might be affected by previous expression of Ino1, leading to a reactivation-specific delay that obscures the effect of memory.

In the presence of inositol, *INO1* transcription is repressed by the combined action of the repressors Opi1 and Ume6 (Graves and Henry, 2000; Jackson and Lopes, 1996; White et al., 1991); loss of either of these proteins leads to constitutive, high-level expression of *INO1*. Depletion of inositol from the medium slows the rate of phosphatidylinositol biosynthesis, leading to an accumulation of the precursor, phosphatidic acid, which directly binds to and inhibits Opi1 (Loewen et al., 2004). Inhibition of Opi1 leads to its dissociation from the promoter, export from the nucleus and activation of *INO1* (Brickner and Walter, 2004; Loewen et al., 2004). In cells that have recently expressed Ino1, the extracellular inositol and the inositol produced by Ino1 may exceed that in cells that have not recently expressed Ino1, delaying the accumulation of phosphatidic acid. To avoid this possible complication, we induced *INO1* transcription by removing Opi1 from the nucleus using the Anchor-Away method (Haruki et al., 2008); Figure 1C). This approach should bypass any insensitivity to inositol starvation, allowing us to directly compare the rate of activation to the rate of reactivation. Upon Anchor Away of Opi1, reactivation was faster than activation and this effect was lost in both *mrs* and *nup100Δ* mutant cells, confirming that it requires the interaction with the NPC (Figure 1D, middle and right panels). Therefore, *INO1* transcriptional memory enhances the rate of transcriptional induction.

We next asked if faster reactivation promoted competitive fitness by competing pairs of strains with nearly identical plasmids integrated into the genome, differing at a single nucleotide (see Methods). The abundance of each strain was quantified by PCR amplification and Sanger sequencing of a segment encompassing the SNP (either A or C; Figure 1E). Mixing strains in various ratios confirmed that this assay is quantitative and accurate over a large dynamic range (Figure 1F). The assay revealed that, during the initial 3h of inositol starvation, cells with inositol memory are more fit than cells experiencing inositol starvation for the first time (Figure 1G). Importantly, this fitness benefit is dependent upon Nup100 (Figure 1G).

Some forms of memory either require transcription during the initial stimulus or reflect the slow dilution of proteins that are expressed in activating conditions (Kundu and Peterson, 2010; Sood and Brickner, 2017; Zacharioudakis et al., 2007). Other forms of memory do not require transcription (Pascual-Garcia et al., 2020). Therefore, we tested if transcription is required to induce *INO1* memory. First, we determined how long cells need to be starved for inositol to induce *INO1* memory using a confocal microscopy-based live-cell localization assay for memory (D’Urso et al., 2016; Egecioglu et al., 2014). The *INO1* locus was tagged with an array of ∼128 Lac repressor binding sites in a strain expressing LacI-GFP (Robinett et al., 1996; Straight et al., 1996). After different periods of inositol starvation followed by 3h in the presence of inositol, the percentage of cells in which *INO1* colocalized with Pho88-mCherry at the nuclear periphery was quantified (Figure 1 – Supplement 1A). Based on the size of the yeast nucleus, we expect a randomly positioned gene to colocalize with the nuclear envelope in ∼27% of cells (blue dashed line; Brickner et al., 2019; Brickner and Walter, 2004). As little as 10 minutes of inositol starvation resulted in Nup100-dependent peripheral localization 3h after shifting back to +inositol (Figure 1 Supplement 1A). This is well before significant transcription has been induced (Figure 1B). Using a temperature-sensitive mutation in the large subunit of RNAPII (*rpb1-1*; Nonet et al., 1987), we find that inactivating RNAPII during inositol starvation did not affect retention at the nuclear periphery after repression (Figure 1 Supplement 1B). Finally, to test if transcription is required in *cis*, we introduced mutations in the TATA box of *INO1*. This mutation blocks *INO1* transcription and leads to an Ino^-^ phenotype (Figure 1 – Supplement 1C). However, localization of *INO1* at the nuclear periphery under either activating or memory conditions was unaffected by mutation of the TATA box (Figure 1 - Supplement 1D). Therefore, neither transcription of a *trans*-acting factor nor transcription of *INO1* is required for localization at the nuclear periphery during memory. This suggests that *cis*-acting molecular changes associated with the early moments of transcription induce *INO1* transcriptional memory.

*INO1* memory persists through ∼ 4 cell divisions (6-8h; Brickner et al., 2007). This duration could reflect either the dilution and degradation of proteins over time or imperfect fidelity of inheritance following DNA replication. To shed light on this, we asked if memory could be extended significantly by arresting/slowing the cell cycle. Indeed, arresting cells during mitosis with nocodazole extended *INO1* memory beyond 18h (Figure 1 – Supplement 1). While dilution is also inhibited by arresting cell division, it seems unlikely that protein production and dilution explain the duration of memory because transcription during inositol starvation is not required for memory. This supports a model in which the duration of memory is regulated by the number of cell divisions or replication cycles.

### A positive feedback loop promotes *INO1* transcriptional memory

To understand the molecular mechanisms controlling the perpetuation and inheritance of *INO1* memory, we tested how the interaction with the NPC impacts chromatin changes and *vice versa*. Chromatin immunoprecipitation (ChIP) in wild type cells reveals that H3K4me2 is observed under activating and memory conditions (Figure 2A), while H2A.Z upstream of the TSS is only observed during memory (Figure 2B; Light et al., 2010). The Sfl1 TF binds to the *INO1* promoter specifically during memory, requires the MRS DNA zip code and is both necessary and sufficient to induce Nup100-dependent peripheral localization (D’Urso et al., 2016). In strains lacking either Nup100 or Sfl1, H3K4me2 is lost during memory (Figure 2A). Incorporation of H2A.Z during memory also requires Sfl1 (Figure 2B), as has been seen for Nup100 (Light et al., 2010). Thus, the interaction with the NPC stimulates both H3K4me2 and H2A.Z incorporation during *INO1* memory (Figure 2E, left).

**Figure 2.**
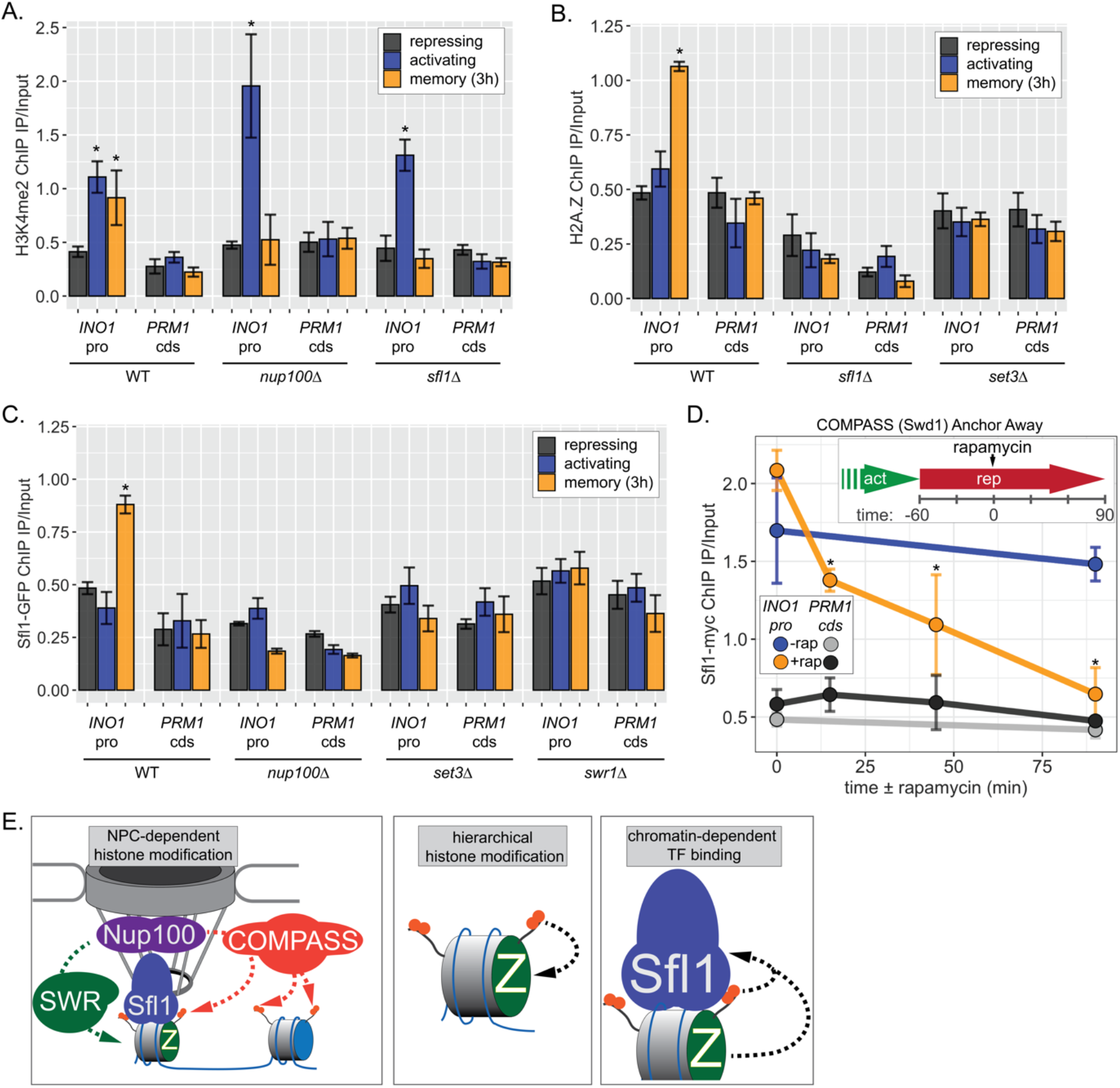
*INO1* transcriptional memory requires a positive feedback loop. **(A - C)** ChIP against H3K4me2 **(A)**, H2A.Z **(B)**, or Sfl1-GFP **(C)** in the indicated strains. Recovery of either the *INO1* promoter or negative control repressed *PRM1* coding sequence was quantified by real-time quantitative PCR and the average of ≥ 3 replicates ± SEM is plotted (*p-value < 0.05 from one-tailed t-test compared with the repressing condition, alternative = greater). **(D)** Time course ± 1µg/ml rapamycin in a Swd1 (COMPASS) Anchor Away strain, quantifying Sfl1-myc ChIP recovery of either the *INO1* promoter or *PRM1* coding sequence under inositol memory conditions (t = 0 is 1h after addition of inositol; *p-value < 0.05 from one-sided t-test compared with the time = 0 minute time point, alternative = less). **(E)** Model of positive feedback relationships: Left: TF-dependent interaction with the NPC stimulates COMPASS-dependent H3K4 dimethylation (orange circles) and SWR-dependent H2A.Z incorporation (green Z). Middle: H3K4me2 is required for H2A.Z incorporation, but H2A.Z incorporation is not required for H3K4me2. Right: H2A.Z and H3K4me2 are required for Sfl1 binding to the MRS.

To explore how chromatin modifications impact each other, we performed ChIP against H2A.Z in a strain lacking Set3, a structural subunit of the SET3C histone deacetylase, which binds H3K4me2 and is essential for maintaining H3K4me2 during memory (D’Urso et al., 2016; Kim and Buratowski, 2009; Light et al., 2013). Strains lacking Set3 failed to incorporate H2A.Z during *INO1* memory, suggesting that H2A.Z incorporation requires Sfl1/Nup100 and H3K4me2. Because H3K4me2 during memory does not require H2A.Z (Light et al., 2013), this suggests a hierarchical relationship between dimethylation of H3K4 and incorporation of H2A.Z (Figure 2E, middle).

Finally, we asked if Sfl1 binding to the *INO1* promoter during memory requires either interaction with the NPC or chromatin modifications. Surprisingly, loss of Nup100, Set3 or Swr1 (the catalytic subunit of the SWR complex, which incorporates H2A.Z into chromatin; Mizuguchi et al., 2004) disrupted binding of Sfl1 to the *INO1* promoter during memory (Figure 2C). In other words, while Sfl1 is required for interaction with the NPC and chromatin modifications, interaction with the NPC and chromatin modification are also required for Sfl1 binding. We confirmed this dependence on H3K4me2 by conditional inactivation of COMPASS (the histone methyltransferase) using Anchor Away of Swd1 after establishing memory (D’Urso et al., 2016). Removing COMPASS from the nucleus leads to loss of H3K4me2 within ∼60min (D’Urso et al., 2016). In cells that have established memory, Sfl1 was bound to the *INO1* promoter (Figure 2D, t = 0). Upon Anchor Away of COMPASS, Sfl1 binding was lost (Figure 2D). Thus, Sfl1 binding to the *INO1* promoter during memory requires H3K4me2 (Figure 2E, right). Together, these data suggest that *INO1* memory involves positive feedback between Sfl1-dependent interaction with the NPC and NPC-dependent H3K4me2 and H2A.Z incorporation.

### Two HSF-like transcription factors are required for *INO1* transcriptional memory

The MRS DNA zip code (5’-TCCTTCTTTCCC-3’; Light et al., 2010) contains sequences reminiscent of the trinucleotide repeats within heat shock elements (5’-TTC-3’). This aided in the identification of Sfl1, which possesses an Hsf1-like DNA binding domain (D’Urso et al., 2016). In budding yeast, there are three other TFs with similar DNA binding domains (Hms2, Mga1 and Skn7), which we also tested for a role in *INO1* memory (Figure 3 – Supplement 1). Of these proteins, only Hms2 was required for localization of *INO1* at the nuclear periphery during memory (Figure 3 – Supplement 1). Loss of Mga1 resulted in constitutive *INO1* localization at the periphery, while loss of Skn7 had no effect (Figure 3 – Supplement 1). This suggests that Sfl1 and Hms2 are specifically required for *INO1* transcriptional memory and that Mga1 may play a role in negatively regulating peripheral localization.

If Hms2 is required for *INO1* memory, we predicted that it would bind to the *INO1* promoter during memory in an MRS-dependent manner and that it would be required for the molecular outputs of memory. ChIP against Hms2-myc revealed that it bound to the *INO1* promoter both in activating and memory conditions and that binding was lost in the *mrs* mutant (Figure 3A). This suggests that Hms2 binds to the MRS both before and after establishing memory. Under activating conditions, loss of Hms2 did not affect RNAPII binding or methylation of H3K4, but during memory, loss of Hms2 led to loss of both (Figure 3B). Therefore, Hms2 binds during both activation and memory but is specifically required for *INO1* memory.

**Figure 3.**
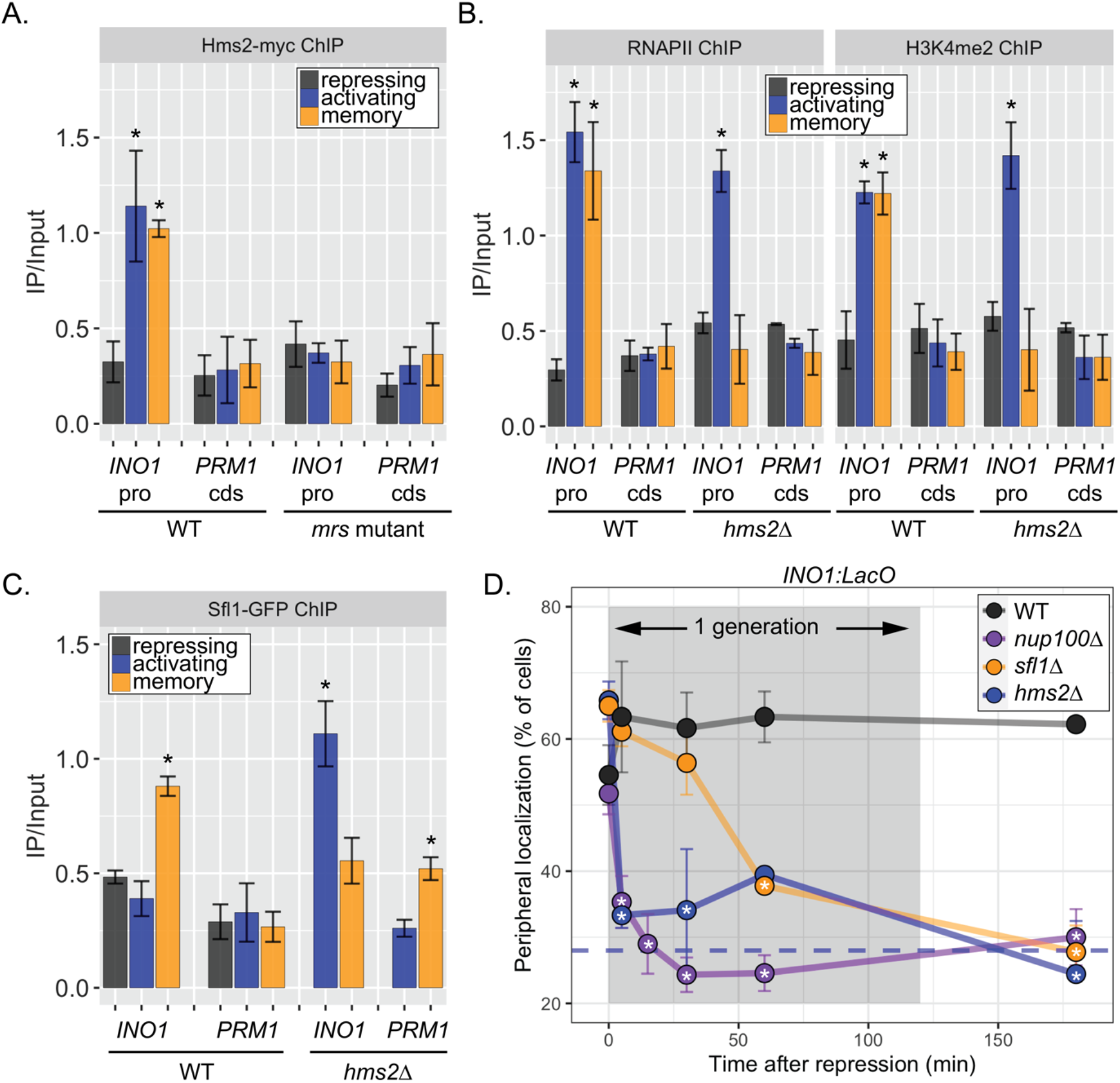
Two different Hsf1-like TFs are required for inositol memory. **(A-C)** ChIP against Hms2-myc (**A**), RNAPII (**B**, left), H3K4me2 (**B**, right) or Sfl1-GFP (**C**) in the indicated strains grown under the indicated conditions. Recovery of either the *INO1* promoter or negative control repressed *PRM1* coding sequence was quantified by real-time quantitative PCR and the average of ≥ 3 replicates ± SEM is plotted (*p-value < 0.05 from one-tailed t-test compared with the repressing condition, alternative = greater). **(D)** Peripheral localization of *INO1* in either wildtype, *sfl1Δ, hms2Δ* or *nup100Δ* strains. At t = 0, inositol was added to cells growing without inositol and peripheral localization was scored at the indicated times. The doubling time of this strain (∼120 minutes) is indicated. The average of three biological replicates ± SEM is plotted and each biological replicate ≥ 30 cells. Blue hatched line: expected peripheral localization for a randomly localized gene. *p-value < 0.05 from one-tailed t-test compared with time = 0, alternative = less.

Because Hms2 bound prior to Sfl1, we tested if Hms2 impacts Sfl1 binding. ChIP against Sfl1-GFP revealed that, in cells lacking Hms2, Sfl1 bound during activating conditions instead of during memory (Figure 3C). Furthermore, loss of either TF or Nup100 led to rapid loss of *INO1* localization at the nuclear periphery, albeit with slightly different kinetics (Figure 3D). All of the mutants resulted in a loss of localization within one generation (denoted by grey box), but loss of Hms2 and Nup100 had the most immediate effect. Thus, our current model is that Hms2 prevents Sfl1 binding to the active *INO1* promoter but is required for Sfl1 binding during memory, perhaps binding as a heterodimer.

### The binding of RNAPII during memory depends on H3K4me2, but H3K4me2 does not depend on RNAPII binding

One of the hallmarks of *INO1* memory is the recruitment of poised RNAPII PIC lacking the Cdk7 kinase (Kin28 in budding yeast) but including the Cdk8 Mediator kinase (Ssn3 in budding yeast; D’Urso et al., 2016). Anchor-Away of Ssn3 from the nucleus disrupts poised RNAPII from the *INO1* promoter during memory, suggesting that Cdk8 is required for recruitment or maintenance of this poised PIC (D’Urso et al., 2016). To test if the kinase activity is required for RNAPII poising during *INO1* memory, we constructed an analog-sensitive allele of Ssn3 (*ssn3-as*; Y236G) that is inhibited by the ATP analog 1-Napthyl-PP1 (NaPP1) (Bishop et al., 2000; Liu et al., 2004). Inhibition of Ssn3 resulted in loss of RNAPII from the *INO1* promoter during memory, without affecting the recruitment of RNAPII under activating conditions (Figure 4A). In contrast, Anchor-Away budding yeast TATA binding protein (Spt15) from the nucleus disrupted RNAPII binding to the *INO1* promoter under both activating and memory conditions (Figure 4A). Inhibition of Ssn3 specifically slowed the rate of reactivation of *INO1* (Figure 4B) and eliminated the fitness benefit or memory (Figure 4D), confirming that poised RNAPII increases the rate of reactivation.

**Figure 4.**
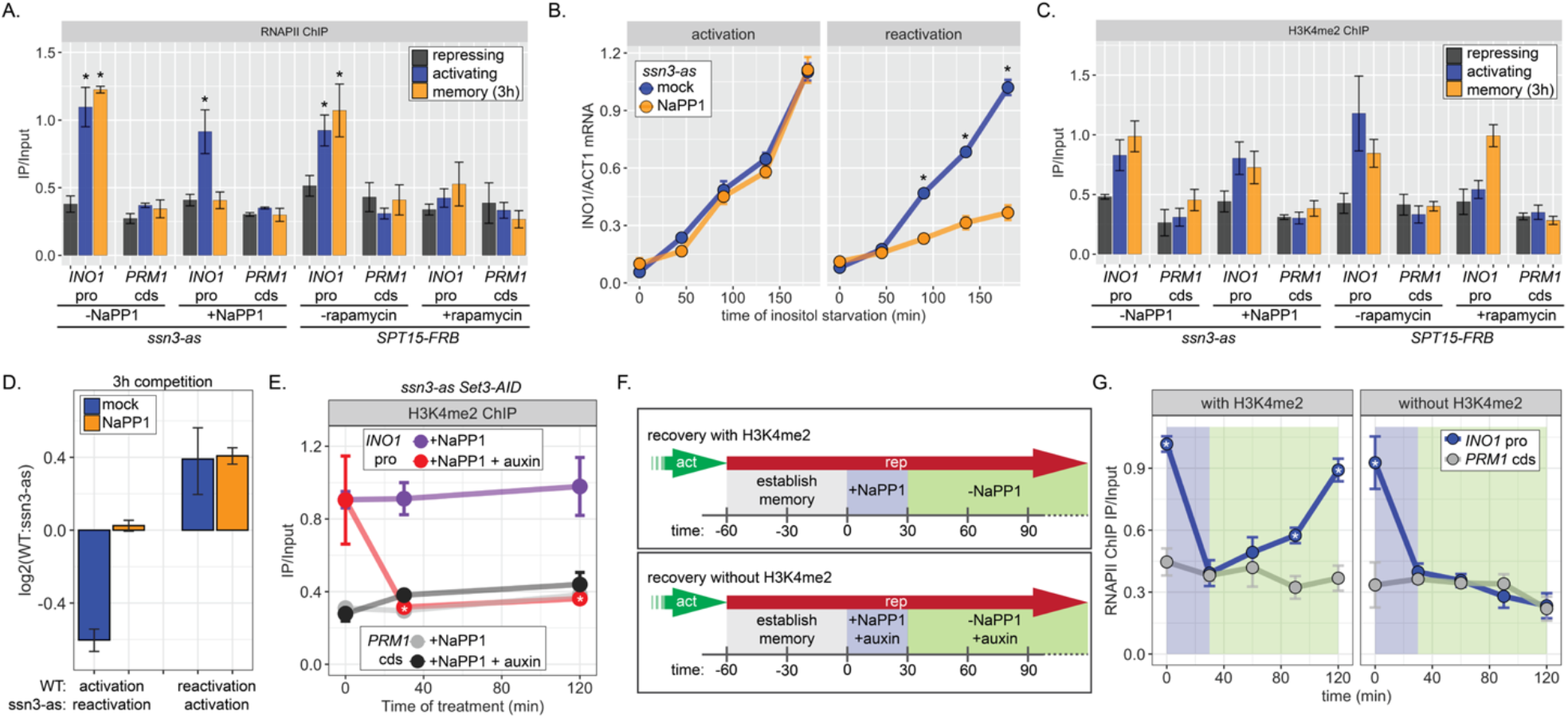
RNAPII binding during *INO1* memory requires H3K4me2, but H3K4 dimethylation does not depend on RNAPII. ChIP against RNAPII **(A)** or H3K4me2 **(C)** in *ssn3* analog-sensitive (*ssn3-as)* and *SPT15-FRB* Anchor Away strains upon addition of either NaPP1 or rapamycin for 1h as indicated. Cells were grown in repressing (+inositol), activating (-inositol), or memory conditions (-inositol ⟶ +inositol, 3 hours). *p < 0.05 from one-tailed t-test compared with repressing condition, alternative = greater. **(B)** Activation (left) and reactivation (right) of *INO1* in the *ssn3-as* strain treated either with DMSO (mock) or NaPP1 for upon inositol starvation (1h pretreatment or treatment). Cells were harvested at indicated time points and the *INO1* mRNA was quantified relative to *ACT1* mRNA by RT-qPCR. *p < 0.05 from one-tailed t-test comparing between mock and NaPP1, alternative = greater. **(D)** Competitive fitness between strains under the indicated conditions. The log_2_ ratio of the strains after 3 hours of competition in media lacking inositol. **(E)** ChIP against H3K4me2 after establishing memory for 1h, followed by addition of either NaPP1 or NaPP1 and auxin at t = 0 (similar to **F** & **G**, but without removing NaPP1). * p-value < 0.05 from one-tailed t-test comparing NaPP1 treated and NaPP1+auxin treated samples at each time, alternative = less. **(F)** Schematic for experiment in **(G)** to monitor RNAPII recruitment with (top) or without (bottom) H3K4me2. NaPP1 was added with or without 0.5mM auxin for 30 minutes before removing NaPP1. **(G)** ChIP against RNAPII, following the experimental set up in **(F)** with cells crosslinked at the indicated times. * p-value < 0.05 from one-tailed t-test comparing ChIP of *INO1* promoter to *PRM1* cds at each time, alternative = greater. For panels **A, C, E** and **G**, recovery of the *INO1* promoter or the *PRM1* coding sequence (negative control locus) was quantified relative to input by qPCR the averages of three biological replicates ± SEM were plotted.

H3K4 methylation is stimulated by active RNAPII (Bae et al., 2020; Krogan et al., 2003, 2002). Consistent with this, Anchor-Away of Spt15 results in loss of H3K4me2 from the active *INO1* promoter (Figure 4C). However, neither depletion of Spt15 nor inhibiting Ssn3 affected H3K4 dimethylation under memory conditions (Figure 4C). This suggests that poised RNAPII at the *INO1* promoter during memory is not required for dimethylation of H3K4.

The inhibition of analog sensitive kinases by NaPP1 is readily reversible (Wan et al., 2006). Therefore, we asked if loss of poised RNAPII from the *INO1* promoter was reversible by testing if RNAPII could return to the promoter upon removal of NaPP1 (Figure 4F, top). Cells were shifted from activating to memory conditions for 1h, Ssn3 was inhibited for 30 minutes to disrupt RNAPII binding and then NaPP1 was washed away (Figure 4F, top). Inhibiting Ssn3 led to loss of RNAPII association with the *INO1* promoter within 30 minutes but, upon removing NaPP1, RNAPII binding was recovered fully within 90 minutes (Figure 4G, left). Therefore, RNAPII is not required to maintain the *INO1* promoter in a memory-compatible state.

To ask if chromatin is critical for the return of poised RNAPII, we asked if RNAPII could return to the *INO1* promoter upon removal of NaPP1 in the absence of H3K4me2 (Figure 4F, bottom). H3K4me2 was removed by auxin-inducible degradation of Set3 (AID; Nishimura et al., 2009), which is required to maintain H3K4me2 over the *INO1* promoter during memory (D’Urso et al., 2016). In the *ssn3-as SET3-AID* strain, adding NaPP1 had no effect on H3K4me2 over the *INO1* promoter during memory, while addition of both NaPP1 and auxin resulted in rapid loss (Figure 4E). In cells treated with auxin, RNAPII failed to return to the *INO1* promoter upon washing away NaPP1 (Figure 4G, right). We conclude that dimethylation of H3K4 is necessary for recruitment of poised RNAPII to the *INO1* promoter during memory and that the pathway for H3K4 dimethylation during memory is independent of RNAPII.

### The Paf1C subunit Leo1 is specifically required for memory

The work on *INO1* suggests that the molecular requirements for dimethylation of H3K4 at active promoters are different from the molecular requirements for dimethylation of H3K4 at the same promoters during memory; while active promoters require RNAPII for H3K4 methylation, poised promoters do not. Therefore, we asked if the Paf1 complex, a conserved factor required for H3K4 methylation, plays a role in H3K4 dimethylation during memory. The yeast Paf1 complex (Cdc73, Ctr9, Leo1, Paf1, and Rtf1), associates with active RNAPII and promotes methylation of H3K4 by recruiting COMPASS (Krogan et al., 2003). Loss of certain proteins (Paf1, Ctr9 and Rtf1) completely blocks H3K4 methylation, while loss of Cdc73 and Leo1 have no obvious effect (Krogan et al., 2003; Ng et al., 2003a, 2003b). We assessed how loss of these factors affected H3K4me2 and H3K4me3 over the *INO1* promoter under repressing, activating and memory conditions. As expected, loss of Ctr9, Paf1, or Rtf1 blocked all methylation and loss of Cdc73 had no effect (Figure 5A & Figure 5 – Supplement 1A). However, strains lacking Leo1 showed normal levels of H3K4me3 and H3K4me2 over the active *INO1* promoter, but no H3K4me2 over the *INO1* promoter during memory (Figure 5A & Figure 5 Supplement 1). Thus, the Leo1 subunit of the Paf1 complex plays a memory-specific role in promoting H3K4 dimethylation.

**Figure 5.**
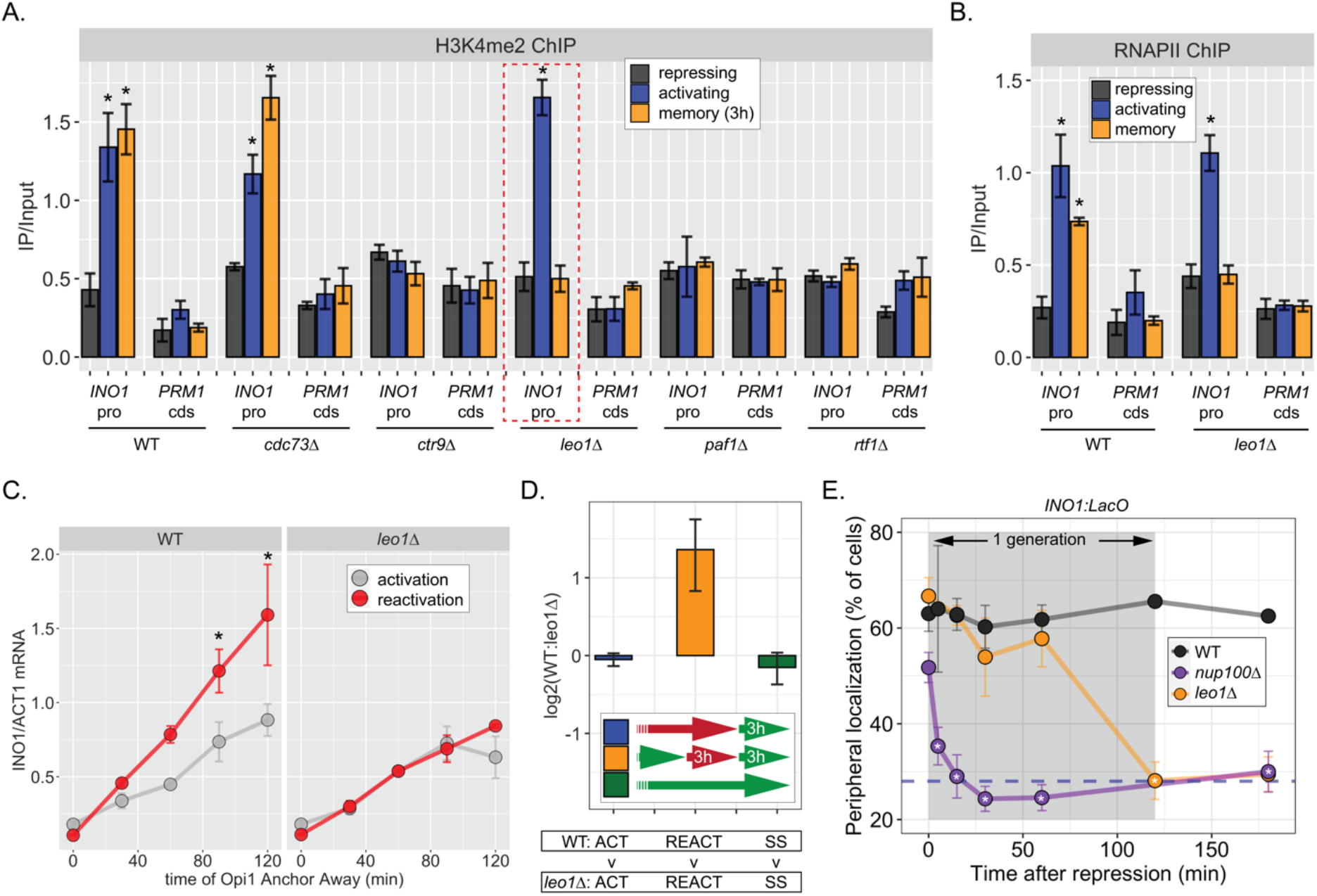
The Leo1 protein of the Paf1 complex is specifically required for *INO1* memory. **(A)** ChIP against H3K4me2 in the indicated strains under repressing (+ino), activating (-ino) or memory (-ino ⟶ +ino, 3h) conditions, highlighting the effects of the *leo1Δ* mutation (dashed red box). **(B)** ChIP against RNAPII in wildtype and *leo1*Δ strains repressing, activating (-inositol) or memory (-inositol ⟶ +inositol, 3h) conditions. For **A** & **B**: recovery of *INO1* promoter or *PRM1* coding sequence (negative control) were quantified by qPCR and the averages of three biological replicates ± SEM are plotted. For A and B, *p < 0.05 from one-tailed t test comparing against repressed condition, alternative = greater. **(C)** Activation and reactivation of *INO1* in wildtype (left) or *leo1Δ* (right) strains upon Anchor Away of Opi1. Cells were harvested at indicated times and *INO1* mRNA was quantified relative to *ACT1* mRNA by RT-qPCR. *p<0.05 from one-tailed t test comparing activation to reactivation, alternative = greater. **(D)** Competitive fitness of wild type vs *leo1Δ* strains competed for 3h in the absence of inositol during activation (ACT, +ino ⟶ -ino, blue), reactivation (REACT, -ino ⟶ +ino (3h) ⟶ -ino, orange), or steady state (SS, -ino ⟶ -ino, green). The ratio of the relative abundance of the WT:*leo1Δ* strains was quantified and expressed as a log2 ratio ± SEM. **(E)** Peripheral localization of *INO1* in wild type, *nup100Δ* (replotted from Figure 3) and *leo1Δ* strains shifted from activating (-ino) to repressing (memory) conditions for the indicated times. The grey box indicates the approximate doubling time of this strain. The average of ≥ 3 biological replicates ± SEM is plotted and each biological replicate ≥ 30 cells.

Consistent with an essential and specific role in *INO1* memory, loss of Leo1 also disrupted *INO1* localization at the nuclear periphery (Figure 5 – Supplement 1B) and RNAPII binding to the *INO1* promoter (Figure 5B) during memory, slowed the rate of transcriptional reactivation (Figure 5C) and erased the fitness benefit associated with *INO1* transcriptional memory (Figure 5D). This essential and specific role of Leo1 in promoting *INO1* memory supports the idea that the molecular mechanism of H3K4 methylation during memory is carried out by an overlapping, but distinct, pathway from that responsible for H3K4 methylation during transcription.

When we examined the rate at which *leo1Δ* lost memory, we found that *INO1* remained at the nuclear periphery for ∼ 60 minutes before repositioning to the nucleoplasm rapidly (Figure 5E). In contrast, loss of Nup100 (or the mutations in the MRS; Light et al., 2010), resulted in immediate loss of peripheral localization upon repression (Figure 5E). Thus, Leo1 is not required to establish memory, but is required to either sustain or inherit memory.

### RNAPII-independent H3K4 dimethylation is mitotically heritable

Because factors such as Leo1 are involved in sustaining, but not establishing memory, we asked if there are factors that play a role in establishing, but not sustaining memory. Sfl1 is required for all aspects of *INO1* memory (D’Urso et al., 2016), Figure 2). To assess if Sfl1 is involved in the establishment of memory, its inheritance, or both, we inactivated Sfl1 using auxin-induced degradation either before or after establishing memory (Figure 6A). Degrading Sfl1-AID either before or after establishing memory led to loss of peripheral localization (Figure 6B), confirming that Sfl1 is critical for interaction with the NPC and that the auxin-induced degradation was complete. Likewise, degrading Sfl1 either before or after establishing memory led to loss of H2A.Z and RNAPII during memory (Figure 1C, left and middle panels). However, H3K4me2 behaved differently. Degrading Sfl1 prior to establishing memory prevented H3K4me2 during memory (Figure 1C, right panel), confirming that Sfl1-AID leads to loss of Sfl1 function. However, in cells in which Sfl1-AID was degraded after establishing memory, H3K4me2 persisted (Figure 1C, right panel). Therefore, once memory has been established, H3K4 dimethylation can persist for up to 2h in the absence of Sfl1.

**Figure 6.**
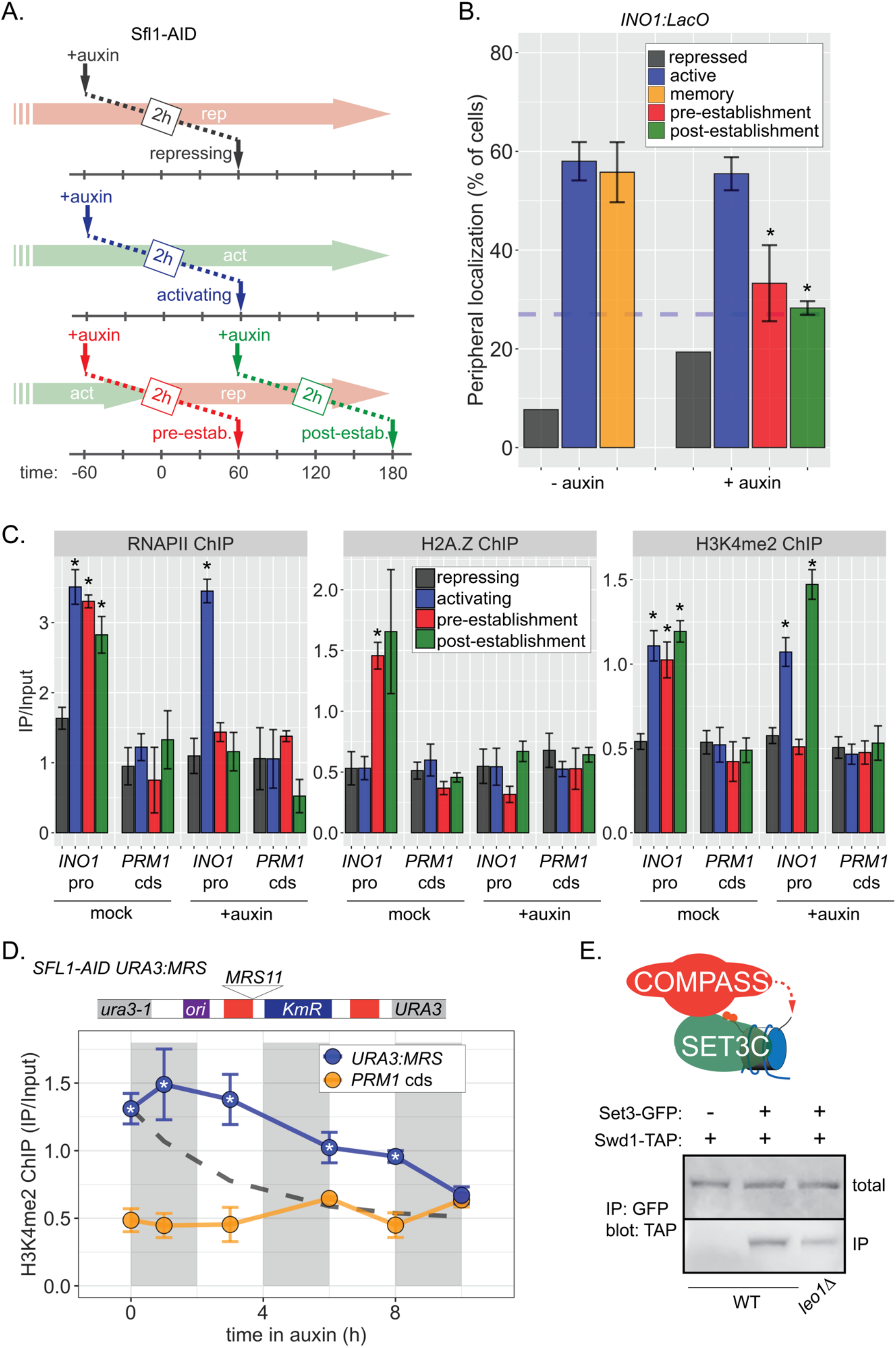
Distinct molecular requirements for establishment and inheritance of H3K4me2 during *INO1* memory. **(A)** Experimental set-up to test the role of Sfl1 in establishment and inheritance of *INO1* memory, using auxin-inducible degradation of Sfl1 before or after establishing memory. Using this approach, peripheral localization of *INO1* (A) or ChIP against RNAPII (**C**, left), H2A.Z (**C**, middle), or H3K4me2 (**C**, right) was measured under the indicated conditions ± 0.5mM auxin. Peripheral localization is the average of three biological replicates ± SEM; each biological replicate ≥ 30 cells. Blue hatched line: expected peripheral localization for a randomly localized gene. *p < 0.05 from one-tailed t-test comparing auxin treated to untreated, alternative = less. **(D)** Schematic of insertion of 11 bp MRS at the *URA3* locus in the *URA3:MRS* strain (top) and ChIP against H3K4me2 in *URA3:MRS Sfl1-AID* strain after addition of auxin (lower). Vertical grey and white bars represent estimated generation times and dashed line represents the expectation from perfect retention of H3K4me2, followed by dilution through DNA replication (*i*.*e*. half-life equal to the doubling time). Panels **C** & **D**: recovery of *INO1* promoter or *PRM1* coding sequence was quantified by qPCR relative to input and are the averages of three biological replicates ± SEM. *p < 0.05 from one-tailed t-test comparing to recovery of each DNA in the repressed condition (**C**) or to *PRM1* cds (**D**), alternative = greater. **(E)** Co-immunoprecipitation of Set3-GFP and Swd1-TAP from the indicated strains. Set3-GFP was immunoprecipitated with anti-GFP nanobodies; recovery of Swd1-TAP was monitored by immunoblotting with anti-TAP antibody.

The observation in Figure 1C raised the possibility that RNAPII-independent H3K4me2 is mitotically heritable in the absence of factors required for its initial deposition. However, to establish the framework for testing this possibility, we first assessed the stability of RNAPII-dependent H3K4 methylation. To do this, we followed H3K4me2 by ChIP over the *mrs* mutant *INO1* promoter following repression, in the absence of transcriptional memory. For comparison, we projected the amount of this mark that would remain if it were neither removed nor deposited (Figure 6 – Supplement 1, dashed line, representing dilution through DNA replication with t½ = 120 minutes). H3K4me2 was lost rapidly over the *mrs* mutant *INO1* promoter upon repression (Figure 6 – Supplement 1), confirming that RNAPII-dependent H3K4 dimethylation is neither stable nor heritable. The active removal of H3K4 methylation upon repression is likely mediated by the H3K4-specific demethylase, Jhd2 (Liang et al., 2007).

To assess if RNAPII-independent H3K4 dimethylation is inherited, we exploited the MRS zip code. Introduction of a single copy of this 11-base pair element near the *URA3* locus (Figure 6D, top) recapitulates important aspects of transcriptional memory: peripheral localization, H3K4me2, and H2A.Z incorporation, but not RNAPII binding (Light et al., 2013, 2010). We have interpreted this to suggest that interaction of the NPC is sufficient to induce the chromatin changes associated with memory, but that recruitment of RNAPII requires *cis* acting promoter elements (D’Urso et al., 2016; Light et al., 2013, 2010). Importantly, the changes induced by the MRS are constitutive, allowing us to assess the duration of heritability without considering the normal mechanisms that regulate the duration of *INO1* memory. Degrading Sfl1-AID led to loss of peripheral localization within 1-2h (not shown), confirming that Sfl1 is required for these effects. However, following Sfl1 degradation, H3K4me2 was maintained for ≥ 8h (∼4 generations; doubling time indicated by the grey and white bars; Figure 6D). This vastly exceeds either the persistence expected from simple dilution of this mark through DNA replication (dashed line) or that observed for RNAPII-dependent H3K4me2 (Figure 6 – Supplement 1). Therefore, RNAPII-independent H3K4 dimethylation is actively reincorporated to allow mitotic inheritance.

The epigenetic maintenance and spreading of histone marks generally requires that the enzyme that catalyzes the deposition of the mark (the writer) physically interact with a protein/complex that recognizes the mark (the reader; Francis, 2009; Ragunathan et al., 2015). Because Set3 both physically interacts with H3K4me2 and is required for transcriptional memory generally (D’Urso et al., 2016), we hypothesized that COMPASS physically interacts with SET3C. Indeed, co-immunoprecipitation of Set3-GFP recovered the COMPASS subunit Swd1-TAP (Figure 6E). This interaction was reduced in a strain lacking Leo1. Therefore, the reader and writer of H3K4 dimethylation physically interact and this interaction is stimulated by Leo1.

## Discussion

Epigenetic phenomena result in heritable changes in phenotype without changes in DNA sequence (Nanney, 1958; Waddington, 2011). Classic examples include stable states, metastable states or regulated switches in state. Stable states include transcriptionally silent subtelomeric, pericentromeric regions or mating type loci (Gartenberg and Smith, 2016; Holoch and Moazed, 2015; Zofall and Grewal, 2006). Metastable states include colony morphology switching in microbes (Klar et al., 2001; Slutsky et al., 1987) and position effect variegation in animals (Elgin and Reuter, 2013). Stable switching of transcriptional states occurs during development, such as silencing of the mammalian X chromosome or establishment of the gene expression programs required for differentiation (Campos et al., 2014; Galupa and Heard, 2018; Gibney and Nolan, 2010; Tee and Reinberg, 2014). Such phenomena often require histone modifications, sometimes called *epigenetic* marks.

However, histone modifications are generally regulated by sequence-specific DNA binding proteins that interact with *cis*-acting, genetically encoded DNA elements (Holoch et al., 2021; Holoch and Moazed, 2015; Laprell et al., 2017). Thus, signal transduction that alters the activity of a TF can produce a long-term change in phenotype that requires histone modifications. Unlike DNA methylation, in which the methyl marks retained on one strand after replication serve to stimulate methylation of the other (Edwards et al., 2017; Hermann et al., 2004), histone/nucleosome modifications are not necessarily re-incorporated at the same location after DNA replication (Escobar et al., 2021; Wang et al., 2009). However, both H3K9 methylation (constitutive heterochromatin; Ragunathan et al., 2015; Zhang et al., 2008) and H3K27 methylation (facultative heterochromatin; Coleman and Struhl, 2017; Hansen and Helin, 2009; Hansen et al., 2008; Margueron et al., 2009) can be inherited through mitosis following removal of the initiating factors (albeit not as stably as with those factors). Furthermore, elegant proximity labeling studies show that nucleosomes at silent loci can be re-incorporated locally after DNA replication (Escobar et al., 2019). Thus, mitotic inheritance reflects both the local reincorporation of marked nucleosomes and reinforcement of these modifications through covalent modification of unmarked nucleosomes following DNA replication. The histone marks for which there is the clearest data for heritability share several features: 1) they tend to occur over large regions, encompassing many nucleosomes, 2) they are associated with transcriptional repression and 3) inheritance requires an interaction between a writer and a reader that recruits or stimulates the writer.

Here we find that H3K4 dimethylation of nucleosomes over the same location in the genome can either be stable and heritable or unstable and rapidly removed, depending on the mechanism by which it is deposited. During active transcription, nucleosomes in the *INO1* promoter are marked with histone acetylation (D’Urso et al., 2016; Rundlett et al., 1998), H3K4me3 and H3K4me2 (Santos-Rosa et al., 2002). This H3K4 methylation is catalyzed by COMPASS and requires RNAPII and the preinitiation complex (Figure 4C; D’Urso et al., 2016). In mutants that lack memory, these marks are rapidly removed upon transcriptional repression (Figure 4 – Supplement 1; D’Urso et al., 2016). In contrast, during memory, histone acetylation and H3K4me3 are lost and H3K4me2 is deposited (Figure 6C; D’Urso et al., 2016). This H3K4me2 is mechanistically distinct from that observed during transcription: it is catalyzed by a subcomplex of Spp1^-^ COMPASS, does not require RNAPII and requires SET3C, Leo1, Sfl1 and Nup100 (D’Urso et al., 2016; Light et al., 2013). Critically, once established, H3K4me2 is both stable and inherited for ∼ 4 cell divisions following degradation of Sfl1 (Figure 6D). This is distinct from the effects of inactivating COMPASS or SET3C (D’Urso et al., 2016). Therefore, H3K4me2 can be maintained and inherited in the absence of an essential initiating factor but continuously requires the writer and reader. We conclude that, as with histone marks associated with heterochromatin (Reinberg and Vales, 2018), a histone mark associated with transcriptional poising is epigenetically inherited.

How is heritable H3K4me2 distinguished from unstable H3K4me2? It is possible that the heritable signal comprises H3K4me2 and additional histone marks. Although the incorporation of H2A.Z during memory is an intriguing possibility for a second signal, it is not essential for H3K4 dimethylation (Light et al., 2013). Alternatively, unacetylated histones may stimulate heritable H3K4 dimethylation, perhaps by regulating the activity of Spp1^-^COMPASS or the Paf complex via Leo1. This may explain why the SET3C histone deacetylase is the reader for this mark.

Whereas H3K9 methylation or H3K27 methylation are generally very stable and state switching promotes long-term or permanent switches, transcriptional memory persists for shorter timescales, generally between 4 and 14 mitotic divisions. This limited inheritance may reflect a balance between the fitness benefits of memory and its costs in a fluctuating environment. Regardless, our current understanding suggests that the duration of memory relates to limits on inheritance following each round of DNA replication. Whereas H3K9 methylation or H3K27 methylation generally covers tens to hundreds of thousands of base pairs (Barski et al., 2007; DiPiazza et al., 2021; Pauler et al., 2009; Yu et al., 2014), the chromatin changes associated with epigenetic transcriptional memory are more local, covering hundreds of base pairs (D’Urso et al., 2016). Following DNA replication, nucleosomes are randomly segregated into the two daughter molecules (Petryk et al., 2018; Yu et al., 2018). Approximately half of the nucleosomes should retain regulatory marks and the inheritance of those marks requires recognizing these nucleosomes and modifying adjacent, unmarked nucleosomes (Escobar et al., 2021; Loyola et al., 2006). If recognition and/or modification is imperfect, the efficiency of such an inheritance mechanism should increase with the number of nucleosomes (DiPiazza et al., 2021). If so, then the duration of memory could be limited by a combination of the number of modified nucleosomes, the fidelity and efficiency of the reader-writer system and the regulation of the initiating transcription factors.

Our current model for *INO1* transcriptional memory is shown in Figure 7. When active, *INO1* interacts with the NPC through upstream DNA zip codes and the TFs Put3 and Cbf1. Two TFs essential for transcription, Ino2 and Ino4, bind to two UAS_INO_ elements that flank the MRS. These TFs recruit co-activators, including histone acetyltransferases like SAGA. Hms2 binds the MRS but has no known role in localization or transcription of active *INO1*. RNAPII recruits the Paf1 complex and COMPASS, leading to H3K4me3 (Figure 7A).

**Figure 7.**
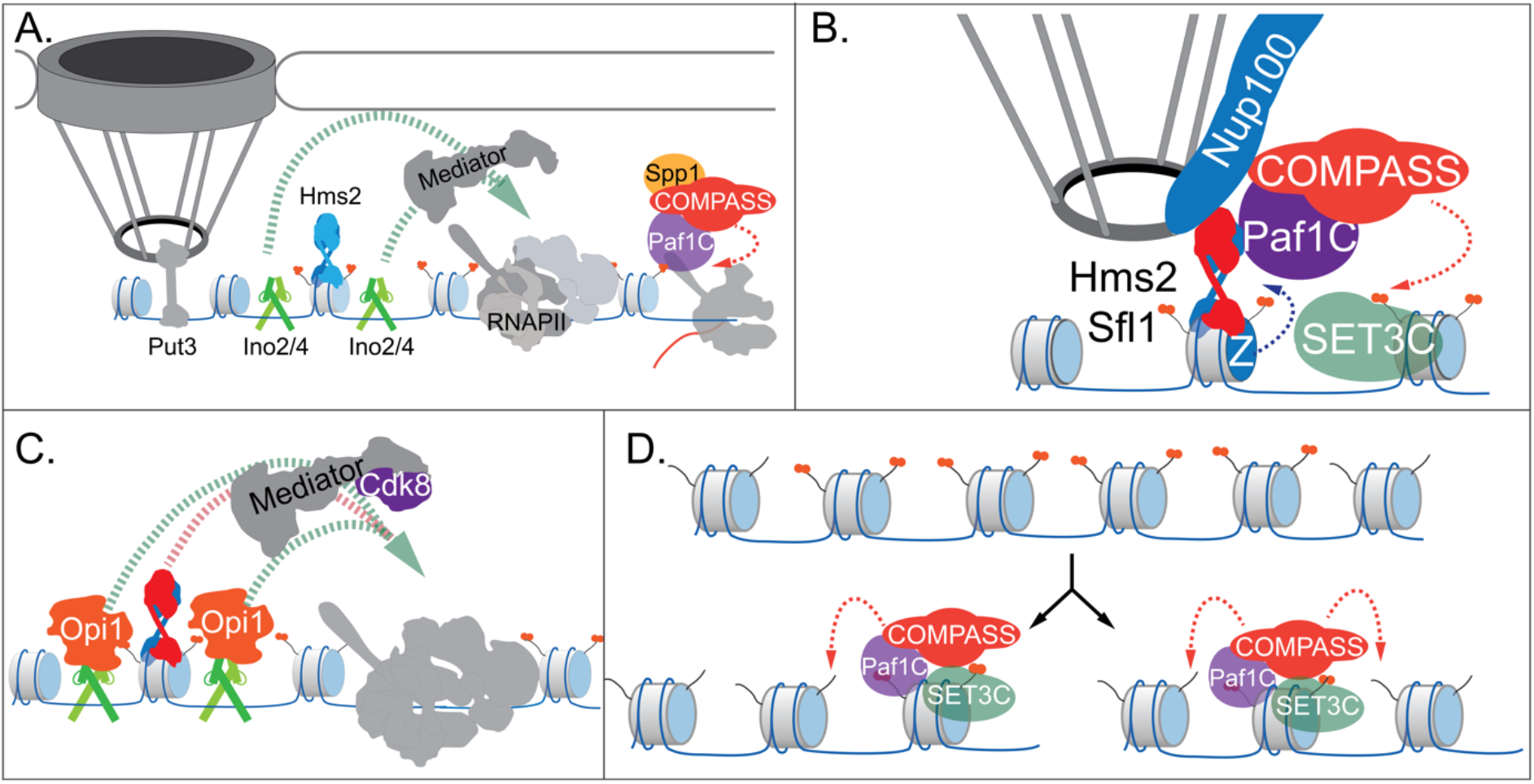
Models for *INO1* transcriptional memory. (**A**). *INO1* under activating conditions. Interaction with the NPC is mediated by Put3 (and, potentially, Cbf1; Ahmed et al., 2010; Randise-Hinchliff et al., 2016). Ino2/Ino4 heterodimers bind UAS_INO_ elements upstream and downstream of the MRS, recruit coactivators such as Mediator to promote RNAPII recruitment and transcription. Hms2 binds to the MRS element, but does not contribute to *INO1* transcription or localization. (**B**). Upon repression, *INO1* transcriptional memory is established by Hms2-dependent recruitment of Sfl1, leading to Nup100-dependent interaction with the NPC. Nup100, Sfl1 and Hms2 are required for both Spp1^-^ COMPASS-dependent H3K4 dimethylation and H2A.Z incorporation near the MRS. SET3C associates with H3K4me2 and is required for its persistence. (**C**) During memory, Cdk8^+^ Mediator is recruited to the *INO1* promoter. This requires Sfl1/Hms2 but may also require other TFs such as Ino2 and Ino4, despite their repression by Opi1. (**D**) H3K4me2 inheritance after DNA replication. Following DNA replication, half of the nucleosomes bear H3K4me2 and the other half do not. SET3C recognizes the H3K4me2-marked nucleosomes and recruits Spp1^-^ COMPASS, which methylates adjacent nucleosomes. In the presence of Sfl1/Hms2, this re-establishment will likely emanate from the MRS outwards.

Upon repression, the Opi1 repressor binds to Ino2 (Brickner and Walter, 2004; Heyken et al., 2005), repressing transcription by recruiting the Rpd3L histone deacetylase (Kadosh and Struhl, 1997; Randise-Hinchliff et al., 2016). To establish memory, Hms2 facilitates recruitment of Sfl1 to the MRS and Nup100-dependent interaction with the NPC (Figure 7B). Because Nup98 physically interacts with the H3K4 methyltransferases Trithorax in flies (Pascual-Garcia et al., 2014) and MLL in mammals (Franks et al., 2017), we envision that Nup100 helps recruit Spp1^-^ COMPASS to dimethylate H3K4 and establish memory (Figure 7B). Perhaps Sfl1, Hms2 or Nup100 compete with Spp1 for binding to COMPASS to ensure this switch. Alternatively, these proteins may recruit the Paf1 complex and COMPASS through interaction with Leo1.

Importantly, the H3K4me2 mark is required for both Sfl1 binding and H2A.Z incorporation during memory, forming a positive feedback loop between interaction with the NPC and chromatin changes. Because the MRS at an ectopic site is sufficient to induce chromatin changes without recruiting RNAPII (Light et al., 2013) or Mediator (our unpublished results), *cis-*acting promoter elements presumably facilitate RNAPII recruitment. We hypothesize that Ino2/Ino4 collaborate with Sfl1/Hms2 to recruit Cdk8^+^ Mediator, which is essential for RNAPII poising (Figure 7C). This would explain why loss of H3K4 methylation leads to loss of Sfl1 binding, peripheral localization and RNAPII.

Following DNA replication, we proposed that inheritance requires methylation of unmodified H3K4 on nucleosomes near those that were previously modified (Figure 7D; Moazed, 2011). This pathway would involve recognition of the H3K4me2 mark on reincorporated nucleosomes by SET3C, recruitment of Spp1^-^ COMPASS and dimethylation of H3K4 on adjacent nucleosomes. Once the *INO1* promoter nucleosomes are methylated, Sfl1/Hms2 can bind, mediating interaction with the NPC and re-establishing memory. Loss of Leo1 seems to both weaken the interaction between COMPASS and SET3C and to disrupt persistence/inheritance of *INO1* memory, suggesting that it facilitates Spp1^-^ COMPASS recruitment after DNA replication.

Do all genes that exhibit transcriptional memory utilize the same molecular mechanism? No. Several of the molecules required for *INO1* memory are not generally involved in memory, such as Sfl1 and Hms2 (our unpublished data). However, *INO1* epigenetic memory has also identified factors and mechanisms that are implicated in memory more generally and suggest a core, conserved, epigenetic poising mechanism. For example, Nup100-dependent interaction with the NPC is associated with transcriptional memory of several yeast genes and Nup98 plays an essential role in both interferon gamma memory in HeLa cells (Light et al., 2010) and in ecdysone memory in flies (Pascual-Garcia et al., 2020, 2017). In yeast and mammals, memory leads to H3K4me2, RNAPII binding and Cdk8 association (D’Urso et al., 2016; Light et al., 2013). Therefore, many genes from yeast to mammals employ a common transcriptional poising mechanism. What selects which genes exhibit memory? Many yeast TFs can mediate Nup100-dependent interaction with the NPC (Brickner et al., 2019). Therefore, different stimuli may induce memory through regulating the activity of these TFs to induce memory in different subsets of genes. Diverse, transient signals could be interpreted through TFs to alter future fitness in distinct ways for several generations.

## METHODS

### Yeast Strains

Yeast strains used in this study are listed in Table 1. Strains used in Co-IP were built using the GFP Library and TAP tag library from Open Biosystems. The *mrs* mutations used in Figure 1 and Figure 4 were created in the endogenous locus via homologous recombination as previously described (Light et al., 2010). Competition strains were created by making SNP mutations in integrative plasmids using inverse PCR (see below). Insertion of MRS11 at the *URA3* locus as found in Figure 7 was done as previously described (Light et al., 2010).

**Table 1.**
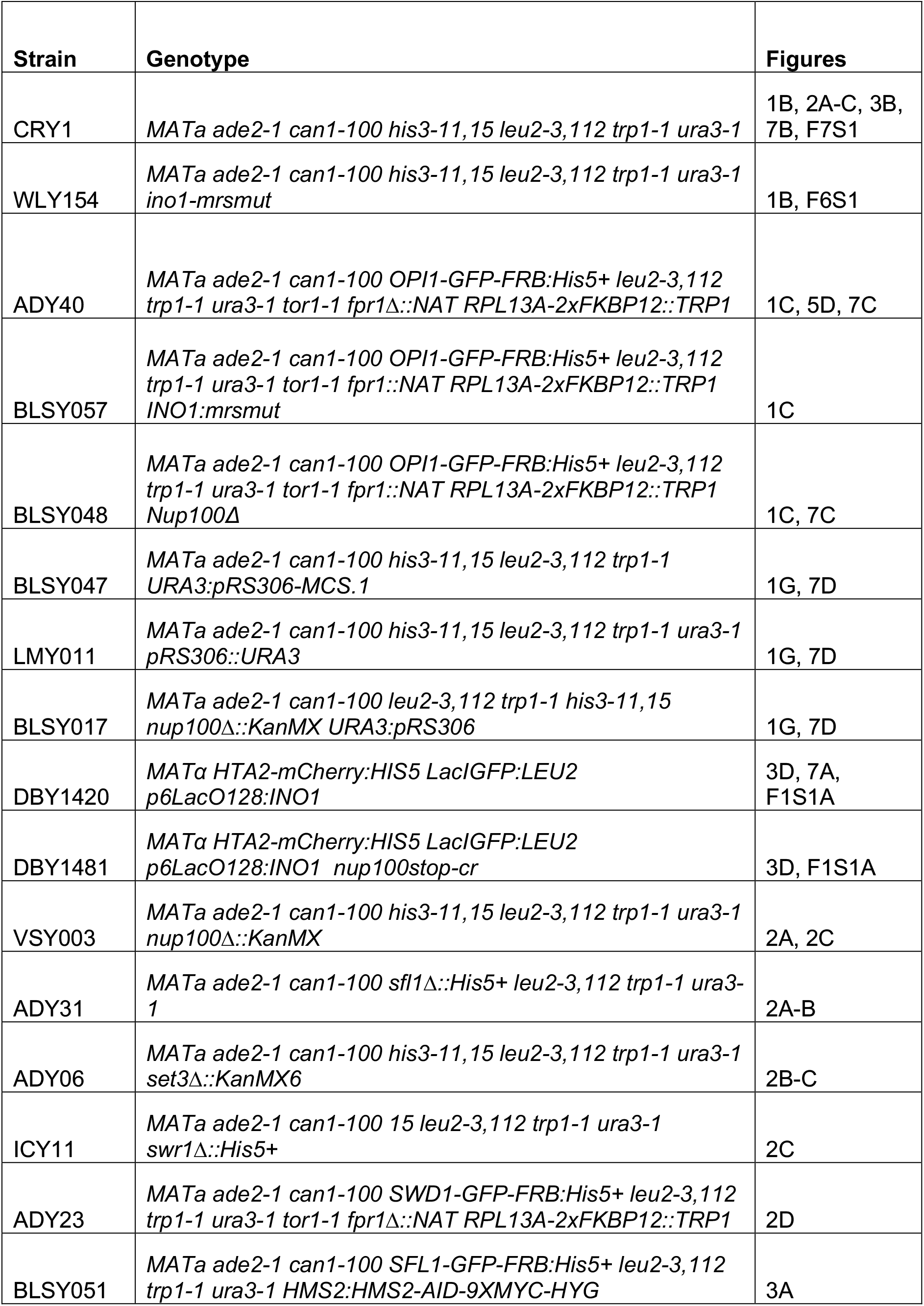
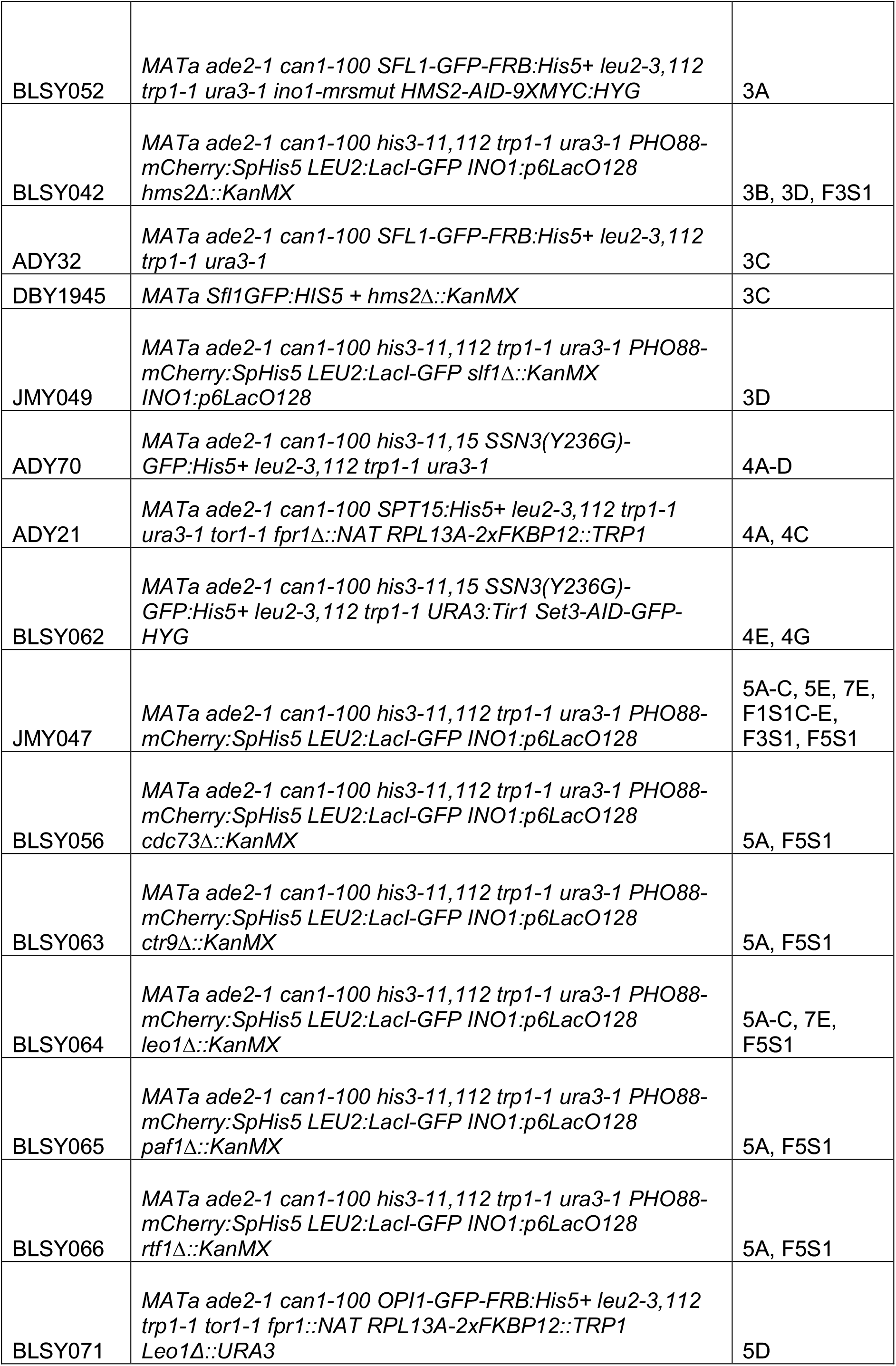
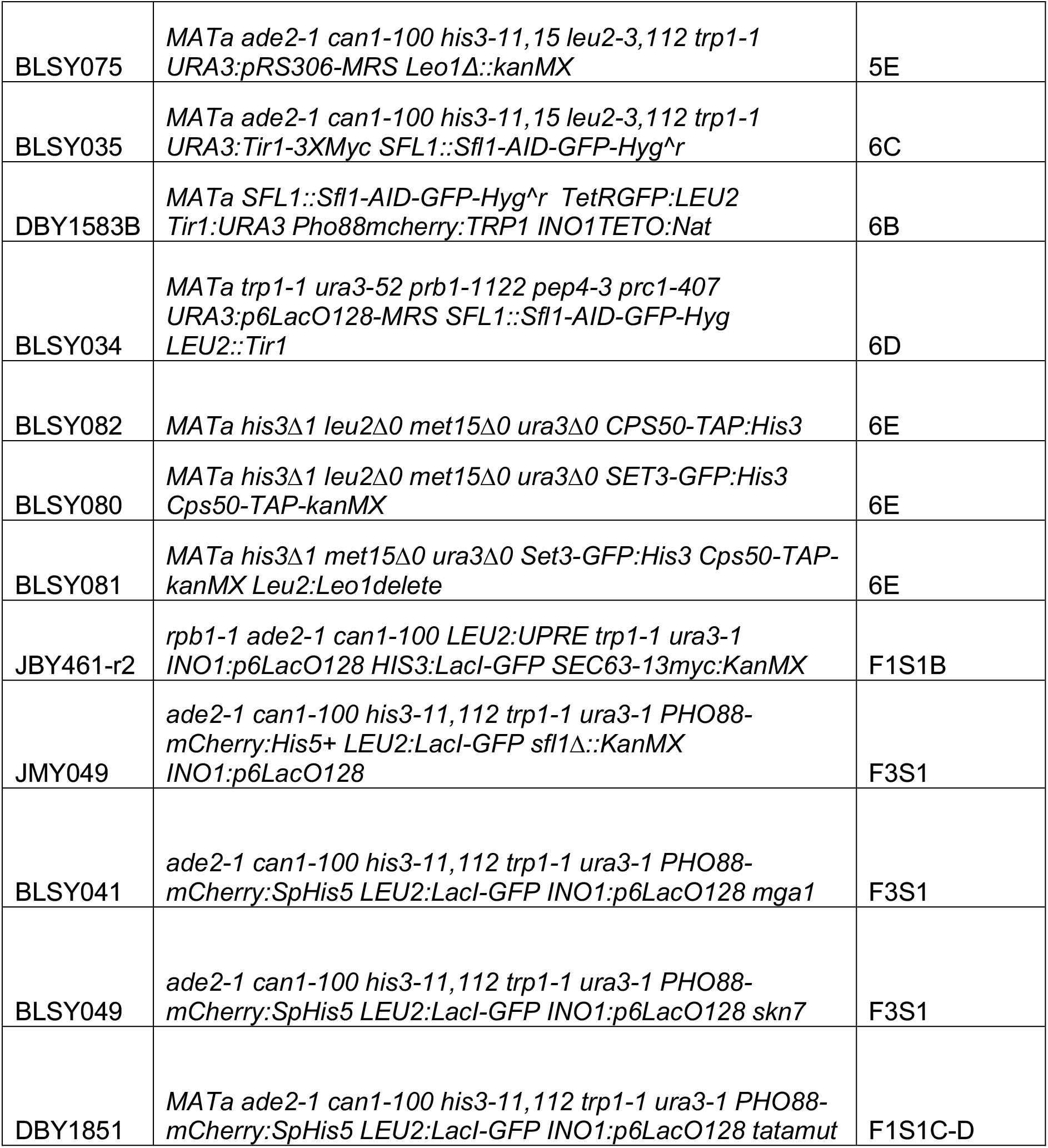
Yeast strains

### Chemicals and Reagents

All chemicals, except those noted otherwise were obtained from Sigma Aldrich (St. Louis, MO). Yeast growth media components were from Sunrise Science (Knoxville, TN) and Genesee Scientific (San Diego, CA). Oligonucleotides (listed in Table 2) are from Integrated DNA Technologies (Skokie, IL), and restriction enzymes from New England Biolabs (Woburn, MA). Antibodies used in ChIP experiments: M-280 Sheep α-Mouse IgG and M-280 Sheep α-Rabbit IgG, from Thermo Fisher Scientific; α -H3K4me2 (ab32356), α-GFP (ab290), α-Myc (ab32), α-H2A.Z (ab4174) from Abcam; α-RNA Polymerase II (cat:664906) from BioLegend; α-TAP (Product#CAB1001) from Invitrogen. The α-GFP nanobody plasmid was a generous gift from Professor Michael Rout (Rockefeller University). The nanobody was expressed, purified and conjugated to magnetic Dynabeads as described (Fridy et al., 2014). The α-H3K4me3 antibody for ChIP was a generous gift from Dr. Ali Shilatifard (Northwestern University).

**Table 2.**
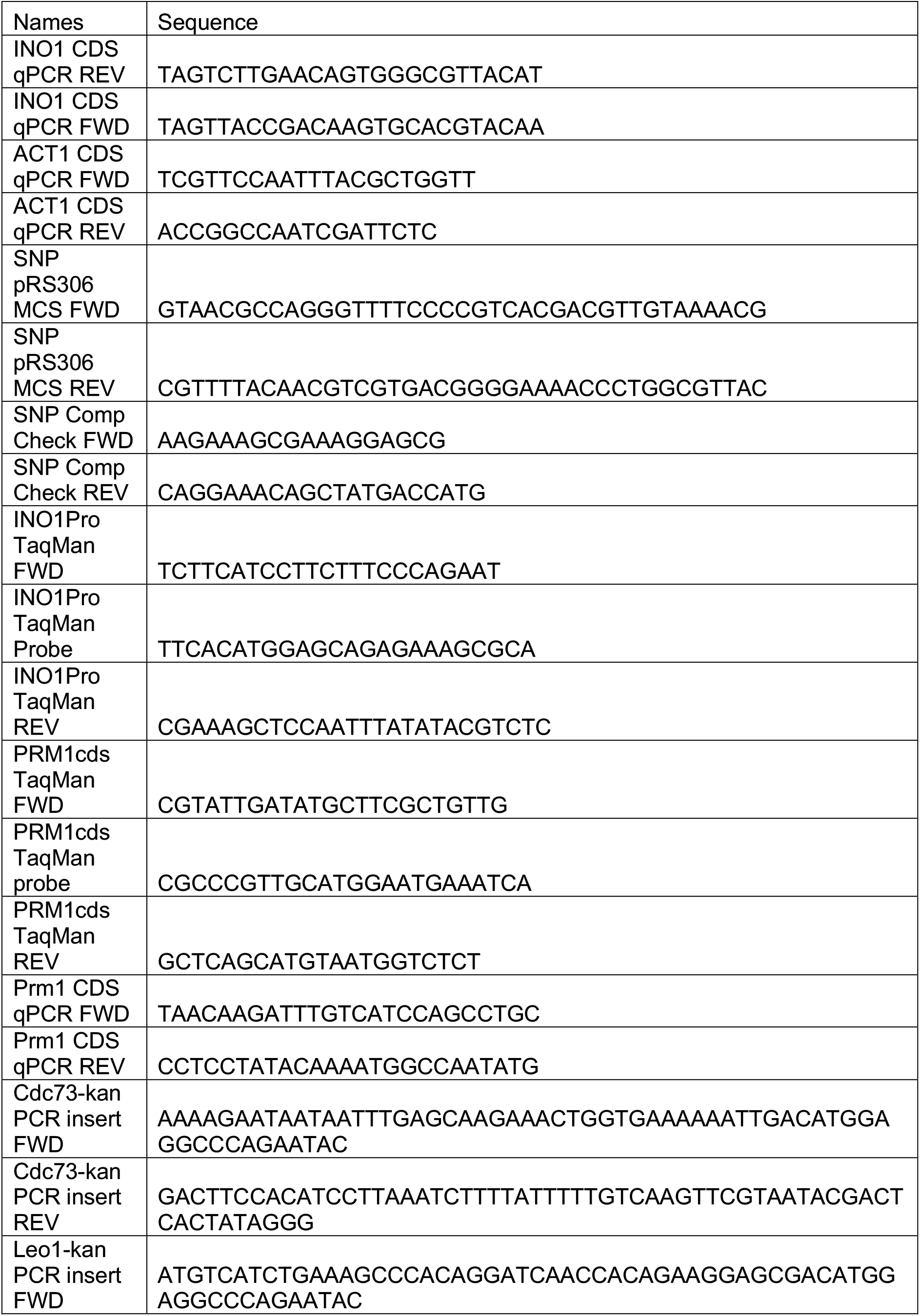
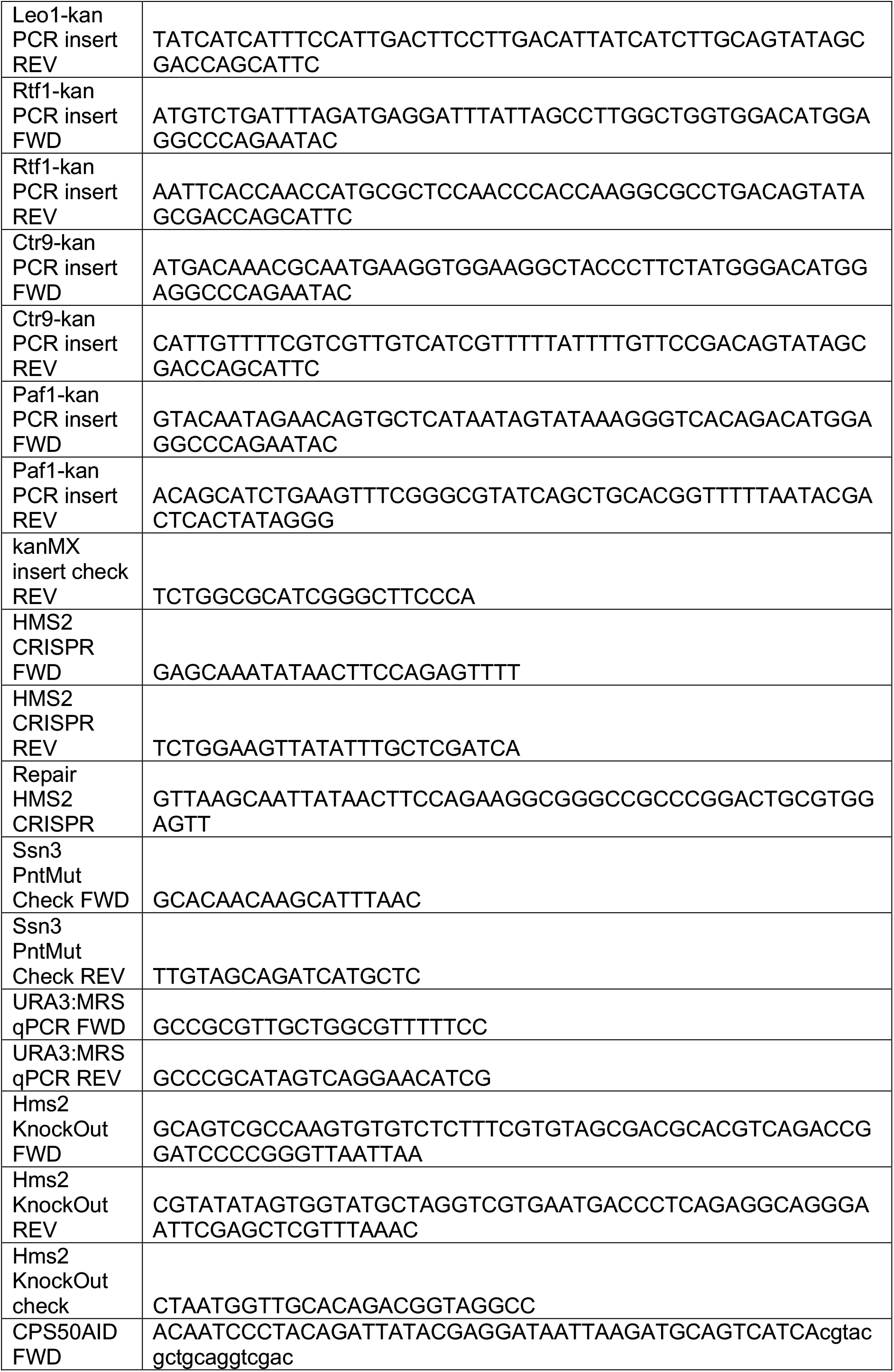
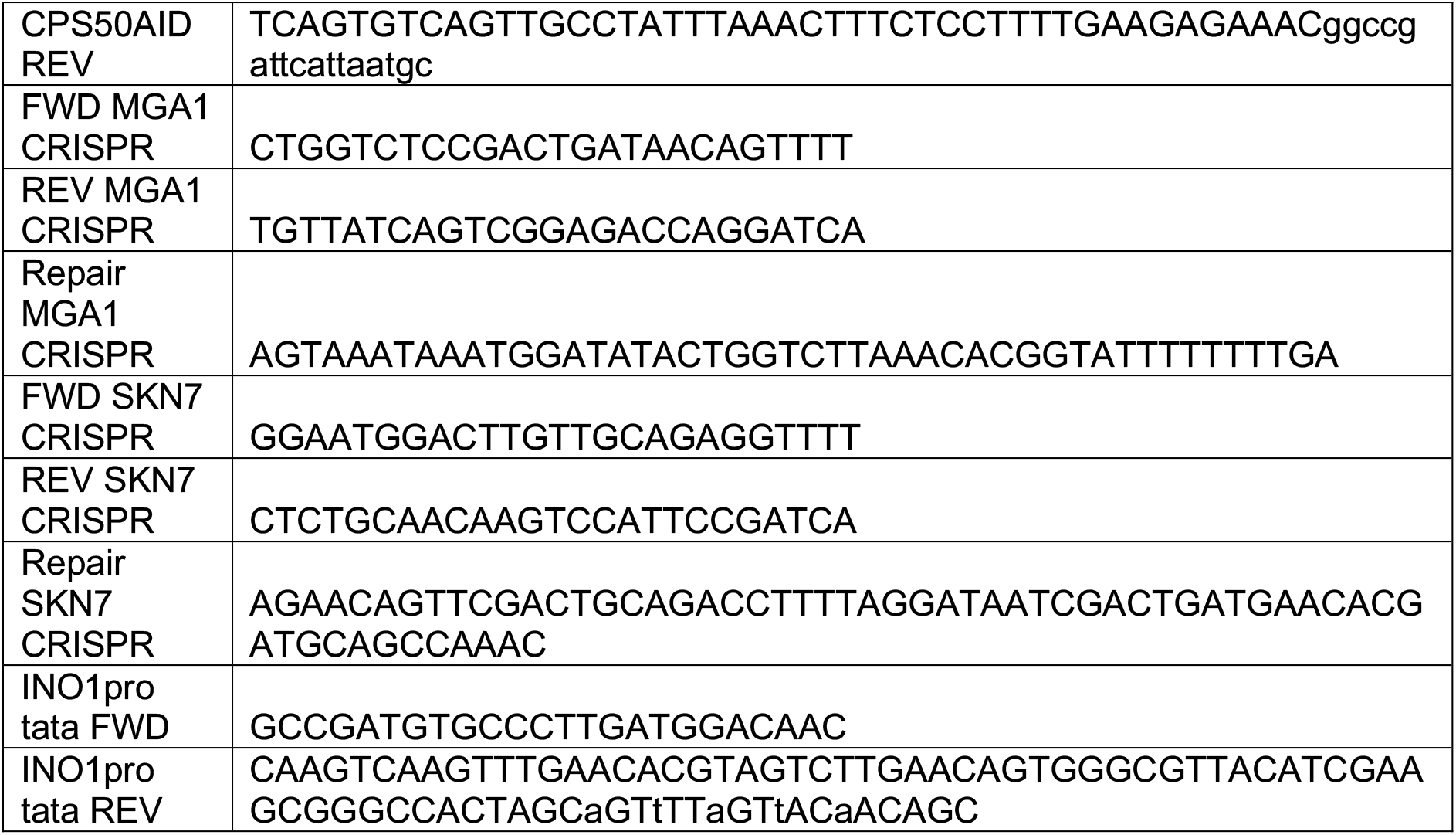
Oligonucleotides

### Chromatin Localization Assay

Chromatin localization was performed as described (D’Urso et al., 2016; Egecioglu et al., 2014), using the confocal SP8 microscope in the Northwestern University Biological Imaging Facility. Error bars represent the SEM of three biological replicates of ≥ 30 cells.

### Reverse Transcriptase Real Time quantitative PCR

For experiments in which mRNA levels were quantified, RT-qPCR was performed as described (Brickner et al., 2007). cDNA recovered from RT was analyzed with traditional qPCR using primers found in Table 2. Error bars represent the SEM of three biological replicates.

### Sequencing-Based Competition Assay

Strains for competition were created by making SNP mutations in the pRS306 (Sikorski and Hieter, 1989) integrating plasmid by inverse PCR using primers found in Table 2. For Figure 1F, the two strains (one carry the SNP mutation “C” and one carrying the wildtype “A” base pair), were combined at various ratios (“A” being 0%, 5%, 25%, 40%, 50%, 60%, 75%, 90%, 100% of the total), estimated using O.D._600_. Genomic DNA was prepared from mixed populations as described (Zentner et al., 2015) and PCR amplified around the SNP region, primers for which are found in Table 2. The resulting chromatograms were analyzed by quantifying the area under the curve of the A/C SNP peak by a custom R script. In the competition experiments, much of the same techniques were used, however the strains would be combined at a 1:1 ratio at the start of the competition (as measured by O.D._600_) and the resulting peak ratio values would be used to normalize the change in peak levels over the 3 h competition period. Additionally, in Figure 5D, the integrative plasmid was not used as the two SNPs already in the strains which created the *ssn3-as* mutation (Y236G) were sufficient to see the difference between the strains after PCR amplification of the genomic preps. The primers used in that panel can be found in Table 2. Error bars throughout represent the SEM of at least three biological replicates.

### Chromatin Immunoprecipitation

ChIP was performed as previously described (D’Urso et al., 2016; Egecioglu et al., 2014). Primary rabbit antibodies (anti-H3K4me2, anti-H3K4me3, anti-GFP, anti-H2A.Z) were recovered with Sheep anti-Rabbit magnetic bead antibodies. Primary mouse antibodies (anti-RNAPII, anti-Myc) were recovered with Sheep anti-Mouse antibodies. DNA recovered from ChIP experiments was analyzed by qPCR using primers in Table 2, using TaqMan qPCR for *INO1* promoter or *PRM1* coding sequence. Error bars represent the SEM of three biological replicates.

### Co-Immunoprecipitation

Co-Immunoprecipitation was performed as previously described (Gerace and Moazed, 2014). When using the GFP nanobodies (described above) to pull down the Set3-GFP, 2 mg/mL of proteins was used as the normalization amount per sample. The Input and IP fractions (after 3 three washes and 10 minutes at 65°C in SDS sample buffer) was analyzed by immunoblotting using rabbit anti-TAP primary antibodies and Goat anti-Rabbit HRP secondary antibodies.

### Immunoblotting

Samples were separated on 10% NuPAGE Bis-Tris gels in MES Running Buffer (Invitrogen), transferred to nitrocellulose, analyzed for total protein loaded with Ponceau stain (Boston BioProducts), blocked with 2% milk for 30 minutes, and incubated with Rabbit anti-TAP primary antibody (Invitrogen) overnight at 4°C. Blots were then washed 3 times with TBS, incubated with Goat anti-Rabbit HRP secondary antibody for 1 hour, washed three times with TBST, then exposed to Enhanced Chemiluminescence reagents (Pierce) for 5 minutes and imaged on an Azure C600 Gel Imaging System.

## Acknowledgements

The authors thank Professor Michael Rout (Rockefeller University) for generously sharing constructs and protocols for the anti-GFP nanobody, Professor Ali Shilatifard (Northwestern University) for generously sharing the anti-H3K4me3 antibody and members of the Brickner laboratory for helpful comments in this manuscript. AD was supported by the Cell and Molecular Basis of Disease T32 training grant (GM008061). This work was also supported by R01 GM118712 (JHB) and R35 GM136419 (JHB).

## Figure legends

**Figure 1 - Supplement 1.**
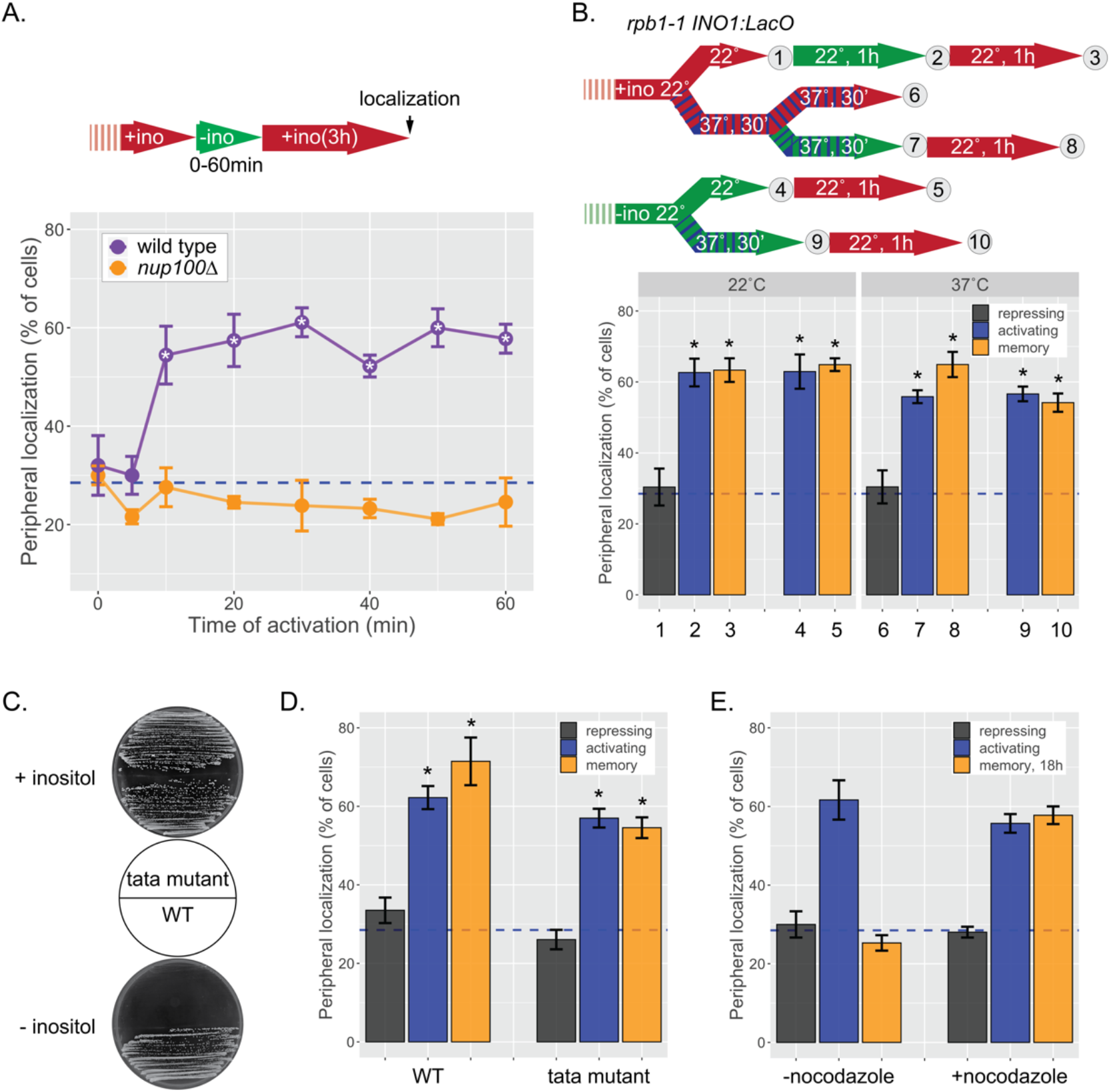
*INO1* memory does not require transcription. **(A)** Cells grown in the repressing (+inositol) condition, switched to the activating conditions (-inositol) for 0-60 minutes, returned to repressing condition for 3 hours before scoring peripheral localization. *p < 0.05 from one-tailed t-test comparing WT to *nup100*, alternative = greater. **(B)** Peripheral localization of *INO1* in a *rpb1-1* temperature sensitive mutant strain. The permissive temperature is 22°C and the restrictive temperature is 37°C. Top: schematic of experimental design; cells were grown under either repressing (red) or activating (green) at either 22°C (solid) or 37°C (hatched) before scoring for peripheral localization. **(C)** Growth of wild type and *tata* mutant cells on SDC +inositol (top) and SDC -inositol (bottom). The TATA box (5’ – TATATAAATT – 3’) at the endogenous *INO1* gene was replaced with 5’ – GCGCGCCCGG -3’ using CRISPR Cas9. **(D)** Peripheral localization of wild type and *tata* mutant *INO1* was scored following 1h of inositol starvation and 3h of growth in +inositol. **(E)** Peripheral localization of *INO1* under repressing (+inositol), activating (-inositol) or memory (-inositol ⟶ + inositol, 18h) conditions, with or without addition of nocodazole for 18h. For all experiments, average of three biological replicates ± SEM; each biological replicate counted a population of ≥ 30 cells. *p < 0.05 from one-tailed t-test comparing to repressed condition, alternative = greater. Blue hatched line: expected peripheral localization for a randomly localized gene.

**Figure 3 - Supplement 1.**
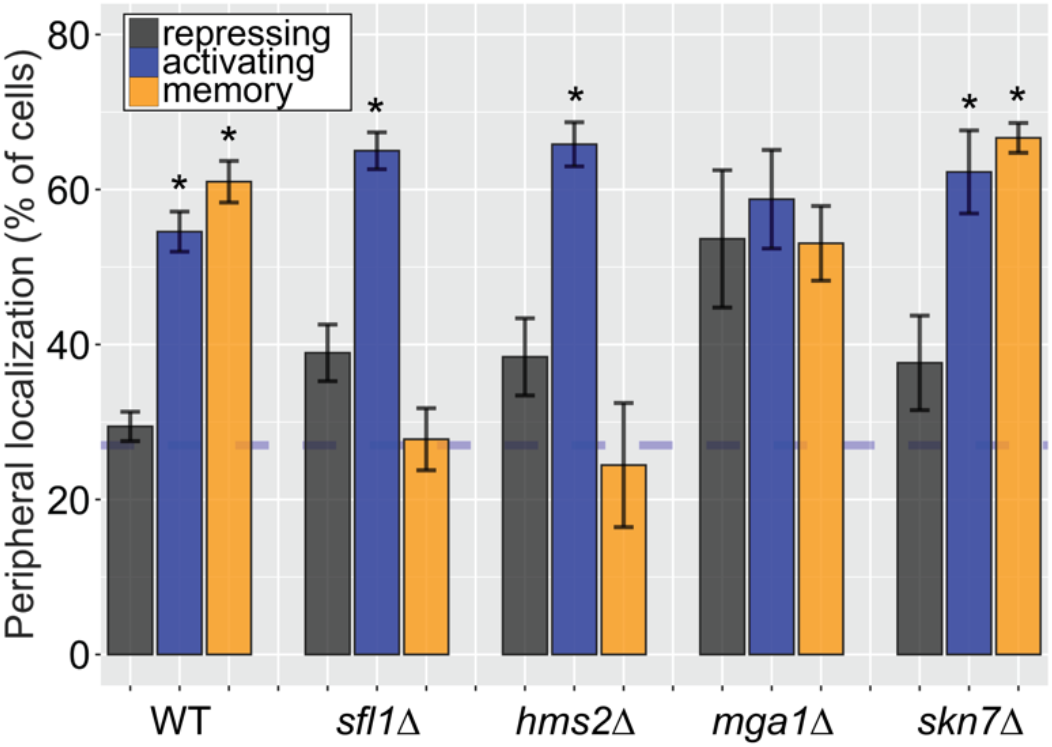
Some, but not all, Hsf1-like TFs are required for *INO1* transcriptional memory. Peripheral localization of *INO1* in either wildtype, *sfl1Δ, hms2Δ, mga1Δ*, and *skn7Δ* strains. Cells grown without inositol were switched to complete media and then imaged after three hours. The average of three biological replicates ± SEM is plotted and each biological replicate ≥ 30 cells. Blue hatched line: expected peripheral localization for a randomly localized gene. *p < 0.05 from one-tailed t-test comparing to the repressing condition, alternative = greater.

**Figure 5 - Supplement 1.**
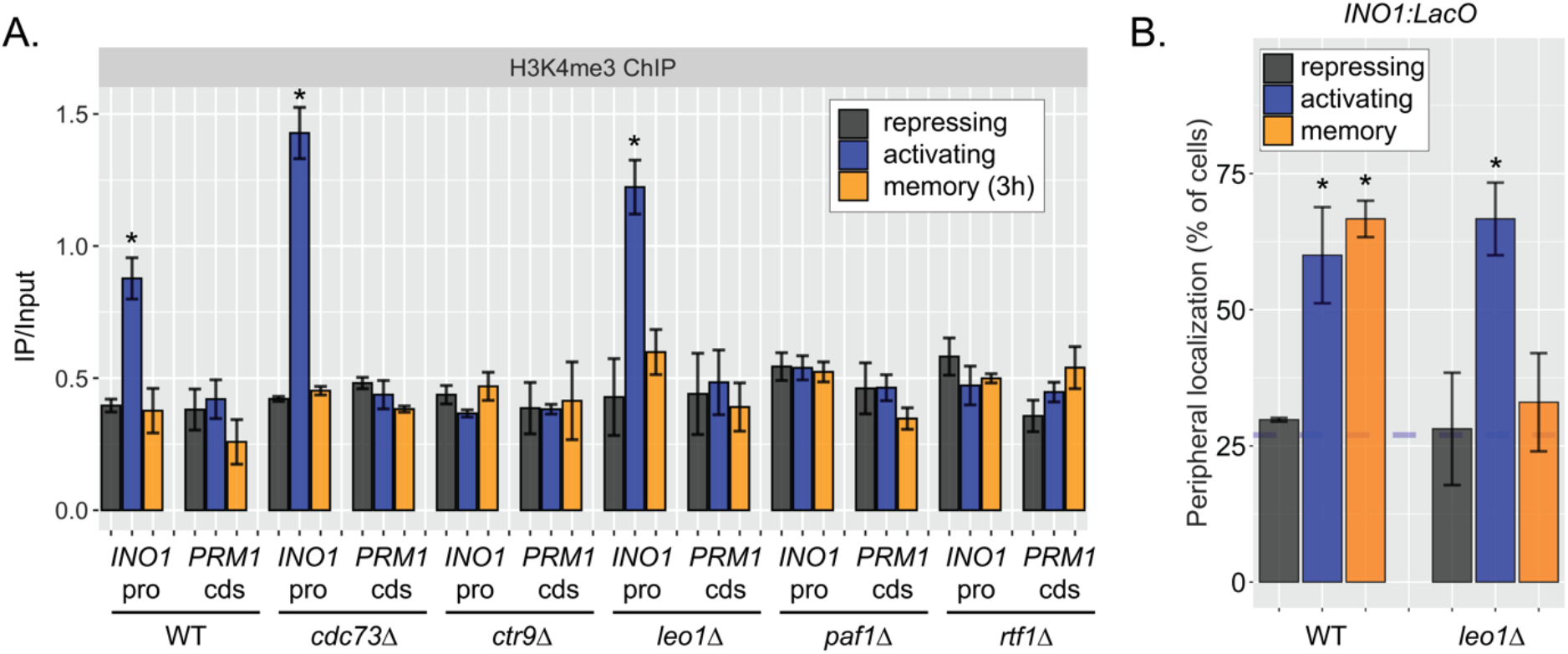
Effects of loss of Leo1 on H3K4me3 and *INO1* localization. **(A)** ChIP against H3K4me3 in wild type, *cdc73Δ, ctr9Δ, leo1Δ, paf1Δ*, and *rtf1Δ* strains in repressing (+inositol), activating (-inositol), memory (-inositol ⟶ +inositol (3h)) conditions. Recovery of the *INO1* promoter or the *PRM1* coding sequence was quantified by qPCR relative to input and are the averages of three biological replicates ± SEM. **(B)** Peripheral localization of wild type and *leo1Δ* strains under repressing (+inositol), activating (-inositol) or memory (-inositol ⟶ +inositol, 3h) conditions. The average of ≥ 3 biological replicates ± SEM; each biological replicate ≥ 30 cells is plotted. *p < 0.05 from one-tailed t test against repressing condition, alternative = greater.

**Figure 6 - Supplement 1.**
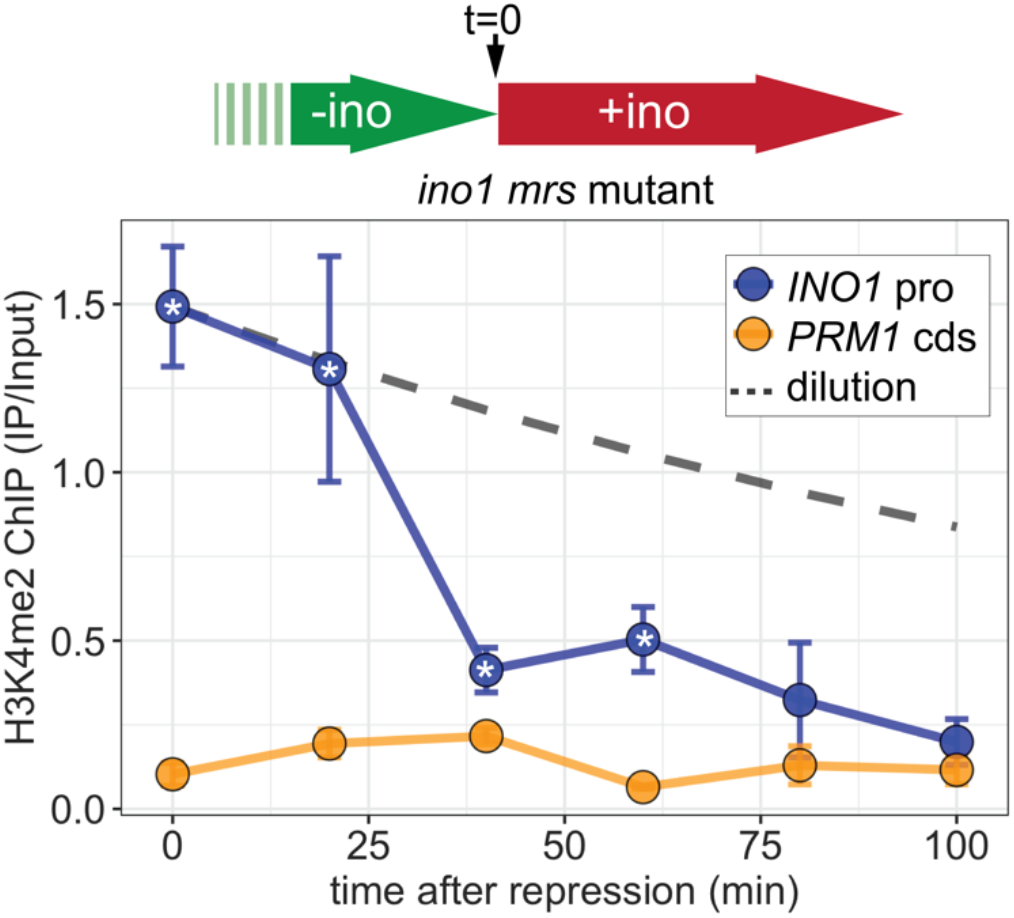
Rapid loss of H3K4me2 from the *INO1* promoter in the absence of transcriptional memory. Time course ChIP against H3K4me2 in the *mrs* mutant strain. Cells were switched from activating (-inositol) to repressing (+inositol) medium at t = 0 and fixed for ChIP at the indicated times. The dashed line represents the expectation from perfect retention of H3K4me2, followed by dilution through DNA replication (i.e. half-life equal to the doubling time). Recovery of *INO1* promoter and *PRM1* coding sequence was quantified by qPCR relative to input and are the averages of three biological replicates ± SEM. *p < 0.05 from one-tailed t-test comparing to recovery of *INO1* promoter to *PRM1* cds, alternative = greater.

